# Distinct evolutionary trajectories following loss of RNA interference in *Cryptococcus neoformans*

**DOI:** 10.1101/2024.08.15.608186

**Authors:** Jun Huang, Connor J. Larmore, Shelby J. Priest, Ziyan Xu, Fred S. Dietrich, Vikas Yadav, Paul M. Magwene, Sheng Sun, Joseph Heitman

## Abstract

While increased mutation rates typically have negative consequences in multicellular organisms, hypermutation can be advantageous for microbes adapting to the environment. Previously, we identified two hypermutator *Cryptococcus neoformans* clinical isolates that rapidly develop drug resistance due to transposition of a retrotransposon, Cnl1. Cnl1-mediated hypermutation is caused by a nonsense mutation in the gene encoding a novel RNAi component, Znf3, combined with a tremendous transposon burden. To elucidate adaptative mechanisms following RNAi loss, two bioinformatic pipelines were developed to identify RNAi loss-of-function mutations in a collection of 387 sequenced *C. neoformans* isolates. Remarkably, several RNAi-loss isolates were identified that are not hypermutators and have not accumulated transposons. To test if these RNAi loss-of-function mutations can cause hypermutation, the mutations were introduced into a non-hypermutator strain with a high transposon burden, which resulted in a hypermutator phenotype. To further investigate if RNAi-loss isolates can become hypermutators, *in vitro* passaging was performed. Although no hypermutators were found in two *C. neoformans* RNAi-loss strains after short-term passage, hypermutation was observed in a passaged *C. deneoformans* strain with increased transposon burden. Consistent with a two-step evolution, when an RNAi-loss isolate was crossed with an isolate containing a high Cnl1 burden, F1 hypermutator progeny inheriting a high transposon burden were identified. In addition to Cnl1 transpositions, insertions of a novel gigantic DNA transposon KDZ1 (∼11 kb), contributed to hypermutation in the progeny. Our results suggest that RNAi loss is relatively common (7/387, ∼1.8%) and enables distinct evolutionary trajectories: hypermutation following transposon accumulation or survival without hypermutation.

**Significance Statement:** There is a dearth of antifungal drugs available to treat *Cryptococcus neoformans*, a human fungal pathogen of global impact. We previously identified natural hypermutators with a loss-of-function mutation in the RNAi machinery and transposon expansion. Here, we identified several novel natural isolates with RNAi defects, none of which are hypermutators or have undergone transposon expansion. Furthermore, we demonstrate that these isolates can lie on a pathway to hypermutation following introduction of a transposon burden. In addition, a novel DNA transposon class was discovered that contributes to antifungal drug resistance. These findings highlight the importance of transposons in driving rapid adaptation in the absence of RNAi and reveal distinct evolutionary trajectories following RNAi loss, a relatively common event in *C. neoformans*.

## Introduction

Stochastic mutations and genomic rearrangements provide genetic variation enabling a microbial population to survive and proliferate in changing environments, driving rapid evolution. However, uncontrolled mutation and rearrangement can be deleterious, leading to gene malfunction and disorganized genome structure. To safeguard the genome, eukaryotic microbes have therefore evolved multiple mechanisms promoting genome stability. Relative to wildtype, microbes with defects in DNA repair, mitochondrial activity, or repressive chromatin modifications have been associated with higher mutation rates, which can lead to distinct adaptability in response to environmental stress (1–6).

RNA interference (RNAi) also serves to promote genome stability (7, 8). Studies in plants, nematodes, and fungi converged to result in the discovery of RNAi, in which the introduction of double-stranded RNA led to the specific silencing of the corresponding gene (9–12). This mechanism plays a critical role in modulating gene expression, defending against viral infection, and maintaining genome integrity (8, 13, 14). As such, RNAi is involved in the formation of heterochromatin, a condensed chromatin structure involved in gene silencing (15). Together with heterochromatin and other cellular mechanisms, RNAi suppresses transposable elements through post-transcriptional or transcriptional silencing within the genome (15, 16).

The RNAi pathway is conserved across a wide range of eukaryotic organisms, but has been independently lost in several fungal species, including the model yeast *Saccharomyces cerevisiae*, the corn smut pathogen *Ustilago maydis*, and the skin commensal/pathogen *Malassezia* species (17–20). As a consequence, several mycoviruses have been found in these RNAi-loss species, which provide benefits by, in some cases, producing toxins (21–23). In addition, shortened centromere length has been correlated with the loss of full-length retrotransposons in the RNAi-deficient *Cryptococcus deuterogattii* species compared to other pathogenic *Cryptococcus* species, highlighting the importance of RNAi in controlling transposons and maintaining genome integrity (24). RNAi was previously thought to be non-functional in the human fungal pathogen *Candida albicans*. However, a recent study found that RNAi is intact in the vast majority of clinical isolates, emphasizing the limitations of utilizing a single reference strain (25).

*Cryptococcus neoformans*, a significant human fungal pathogen in the *Cryptococcus* pathogenic species complex, causes the majority of cryptococcal meningoencephalitis and poses a significant challenge to global human health (26, 27). Population analysis has revealed four distinct *C. neoformans* lineages: VNI, VNII, VNBI, and VNBII (28). Cryptococcal meningoencephalitis is associated with ∼112,000 annual deaths and accounts for ∼19% of all HIV/AIDS-related mortality worldwide (29, 30). Accordingly, *C. neoformans* was classified as a critical priority pathogen in the fungal priority pathogens list published by the World Health Organization (WHO) in 2022 (31, 32).

Unfortunately, options for treating cryptococcal infections are limited. The combination of amphotericin B and 5-fluorocytosine (5-FC) remains the gold standard treatment strategy with improved survival, albeit with side effects (33, 34). However, 5-FC monotherapy is untenable due to rapid development of resistance (35). Similarly, *C. neoformans* resistance to azoles (e.g., fluconazole) that are commonly used for maintenance therapies frequently develops via aneuploidy or mutations (36, 37).

Several natural and laboratory hypermutator isolates have been identified in the *Cryptococcus* pathogenic species complex, with increased mutation rates mediated by a variety of mechanisms. *C. neoformans* and *C. deuterogattii* isolates with mutations in components of the DNA mismatch repair pathway (e.g., *MSH2*) showed elevated mutation rates in homopolymeric tracts, following selection for antifungal drug resistance as well as rapid phenotypic switching (4, 38, 39). Additionally, disruption or mutation in *C. neoformans* of Rad51, a core component of homologous recombination, and in *C. deneoformans* of DNA polymerase delta subunit (*POL3*) are associated with an elevated mutation rate. (2, 40, 41). Furthermore, transposition of the T1 DNA transposon and the TCN12 retrotransposon into the 5-Fluoroorotic Acid (5-FOA) drug targets (*URA3* and *URA5*) underlies the vast majority of 5-FOA resistant colonies isolated from mice infected with the *C. deneoformans* strain XL280α (42).

Recently, we identified two natural hypermutators, Bt65 and Bt81 (43). These strains display a significantly higher mutation rate compared to H99, the type strain for *C. neoformans*, when selected for resistance to a combination of the antifungal drugs rapamycin and FK506 (43). Interestingly, the frequent movement of a non-LTR retrotransposon (i.e., Cnl1) into the gene encoding the FK506/rapamycin target FKBP12 is responsible for most of the resistance. Both hypermutator isolates harbor a nonsense mutation in the gene encoding a novel RNAi component, Znf3, and have accumulated a tremendous Cnl1 transposon burden (43). Further evidence supports the model that the loss of RNAi and high Cnl1 burden are together responsible for hypermutation and antifungal drug resistance (43, 44).

In this study, we sought to determine if other natural *C. neoformans* isolates have lost RNAi and display a similar hypermutator phenotype. Taking advantage of publicly available whole-genome sequencing data for 387 *C. neoformans* natural isolates, we successfully identified five novel RNAi-deficient strains from clinical and environmental sources. Surprisingly, unlike Bt65 and Bt81, none of these RNAi-loss strains display a hypermutator phenotype or contain a Cnl1 transposon burden. We demonstrated that these novel RNAi-deficient strains may lie on a pathway to hypermutation following transposon accumulation. Notably, we discovered several novel gigantic DNA transposons from the KDZ (Kyakuja, Dileera, and Zisupton) class with diverse regulatory mechanisms, contributing to the hypermutator phenotype of RNAi-deficient strains in *C. neoformans* and *C. deneoformans*. Taken together, our results suggest natural isolates with RNAi defects are relatively common and that they may exist at an evolutionary fork in the road, posed to either evolve into hypermutators as transposons accumulate, or persist without such changes despite the loss of RNAi.

## Results

### RNAi loss-of-function mutations occur frequently in *C. neoformans* natural isolates

To broadly analyze genetic variation among natural isolates, two pipelines were developed that utilize short-read sequencing data to identify mutations by mapping reads to a fully assembled and well-annotated reference genome. The first pipeline, Neostop, specifically identifies nonsense mutations, while the other, GenesComp, characterizes all genetic variants, including those occurring within the coding sequences (e.g., synonymous/nonsynonymous substitution, frameshift insertion/deletion, and nonsense mutation), as well as at intron splice sites. Both pipelines were applied to analyze the 387 *C. neoformans* natural isolates in the Strain Diversity Collection using publicly available whole genome sequencing data and the genome assembly and annotation of type strain H99 as reference (28, 45). We focused on genes encoding known components required for the RNAi pathway (SI Appendix, Table S1) (43, 46–50) and on variants predicted to cause loss-of-function (LOF) mutations, namely nonsense, frameshift, and splice-site mutations due to their high probability of causing truncated or dysfunctional gene products.

In total, we identified LOF mutations in five genes required for RNAi in 11 natural isolates (Table 1). Specifically, Neostop and GenesComp detected mutations in six and eight isolates, respectively, with three isolates detected by both pipelines. Most of the isolates had an LOF mutation in only one of the genes analyzed, with the exception of strain LP-RSA2296 which had LOF mutations in two RNAi-related genes *ZNF3* and *AGO1*. The mutations detected included the well-known nonsense mutation in the first exon of *ZNF3* in two hypermutator strains, Bt65 and Bt81, thus, providing a proof-of-principle for this approach (Fig. 1A) (43). All but one of the identified mutations were validated by PCR and Sanger sequencing; strain Bt84 was confirmed to be a false positive and excluded from further analysis (Table 1, SI Appendix, Dataset S1 and Fig. S1). Additionally, the splice-site mutation identified in the *RDE4* gene of strain Bt210 was validated by reverse transcription PCR (RT-PCR) and sequencing of the amplicon encompassing the intron 3/exon 4 boundary, confirming the predicted single nucleotide splicing error at the 3’-end of the third intron, leading to a single nucleotide frame-shift insertion resulting in the introduction of a premature stop codon one amino acid into exon 4 (Fig. 1B and SI Appendix, Fig. S1).

**Figure 1.**
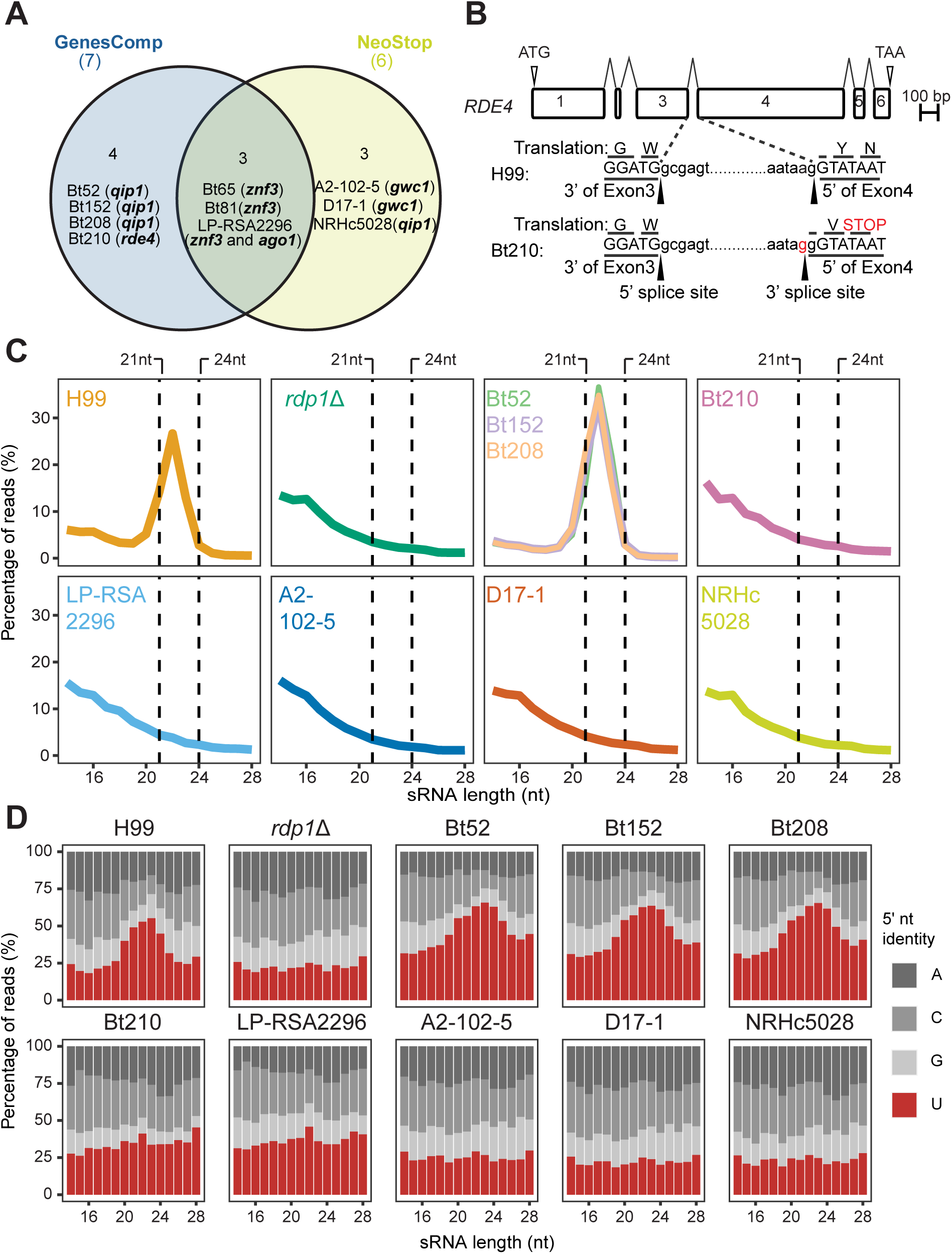
Identification and confirmation of RNAi loss-of-function mutations in *C. neoformans* natural isolates. (A) Venn diagram illustrating the 11 RNAi loss-of-function mutations initially predicted by the two independent pipelines, GenesComp and NeoStop, among *C. neoformans* natural isolates in the Strain Diversity Collection. (B) *RDE4* gene structures in H99 and Bt210 are illustrated. The *RDE4* splice site mutation (A>G) found in Bt210 is highlighted in red. (C) Size distribution of sRNA reads from each of the indicated strain. Dashed vertical lines represent the 21 to 24 nt size range, corresponding to the RNAi-mediated sRNA population. (D) Proportion of 5’-nucleotide identity in sRNA reads. All the sRNA reads were mapped to the H99 genome.

**Table 1.** Natural *C. neoformans* isolates with putative loss-of-function mutations in genes required for RNAi-mediated gene silencing.

| Strain | Pipeline | Gene with mutation <sup>1</sup> | Mutation type | Putative mutation impact <sup>2</sup> |
| --- | --- | --- | --- | --- |
| Bt84 | GenesComp | *QIP1* | Nonsense; Exon 1 | Protein truncated to 30 aa (4% WT length) |
| Bt65 | NeoStop, GenesComp | *ZNF3* | Nonsense; Exon 1 | Protein truncated to 96 aa (6% WT length) |
| Bt81 | NeoStop, GenesComp | *ZNF3* | Nonsense; Exon 1 | Protein truncated to 96 aa (6% WT length) |
| LP-RSA2296 | NeoStop, GenesComp | *ZNF3* | Nonsense; Exon 1 | Protein truncated to 176 aa (11% WT length) |
| Bt52 | GenesComp | *QIP1* | Start-loss; Exon 1 | Missing first 15 aa; truncated to 732 aa (98% WT length) |
| Bt152 | GenesComp | *QIP1* | Start-loss; Exon 1 | Missing first 15 aa; truncated to 732 aa (98% WT length) |
| Bt208 | GenesComp | *QIP1* | Start-loss; Exon 1 | Missing first 15 aa; truncated to 732 aa (98% WT length) |
| LP-RSA2296 | NeoStop, GenesComp | *AGO1* | Frameshift; Exon 10 | Deletion of 9 nt and 1 nt in Exon 10; truncated to 459 aa (50% WT length) |
| Bt210 | GenesComp | *RDE4* | Splice-site; Intron 3 | Splicing error in Intron 3 leads to frameshift; protein truncated to 207 aa (43% WT length) |
| A2-102-5 | NeoStop | *GWC1* | Frameshift; Exon 1 | Protein truncated to 47 aa (8% WT length) |
| D17-1 | NeoStop | *GWC1* | Frameshift; Exon 1 | Protein truncated to 47 aa (8% WT length) |
| NRHc5028.<br>ENR.STOR | NeoStop | *QIP1* | Nonsense; Exon 4 | Protein truncated to 250 aa (33% length) |

<sup>1</sup>: *ZNF3*: CNAG\_02700; *QIP1*: CNAG\_01423; *AGO1*: CNAG\_04609; *RDE4*: CNAG\_01157; *GWC1*, CNAG\_06486
<sup>2</sup>: WT: wild-type; aa: amino acids

Furthermore, the effects of the identified putative RNAi LOF mutations on small RNA production were analyzed by small RNA-sequencing (sRNA-seq), with H99 wild-type and H99 *rdp1*Δ strains serving as RNAi-proficient and -deficient controls, respectively (Fig. 1C and 1D). Our data showed that five of the strains tested (Bt210, LP-RSA2296, A2-102-5, D17-1, and NRHc5028.ENR.STOR (hereafter referred to as NRHc5028)) had sRNA profiles comparable to the RNAi-deficient control H99 *rdp1*Δ, that is, there was neither enrichment of sRNA species in the size range 21-24 nucleotides nor enrichment of 5’-uracil in the small RNA reads. This corresponds well with the size and feature of canonical small inferencing RNAs (siRNAs) utilized in RNAi-mediated silencing, thus confirming that these strains are indeed RNAi-deficient. On the contrary, strains Bt52, Bt152, and Bt208 had small RNA profiles similar to the RNAi-proficient control H99, with enrichment of both 21-24 nt species as well as a preference of 5’-uracil among these small RNAs, suggesting that while these strains have mutations in the *QIP1* gene, they are still able to produce small RNA species characteristic of RNAi-proficient strains (Fig. 1C and 1D). Interestingly, these three strains all have the same start-loss mutation in exon 1, which is predicted to result in a 15-amino-acid truncation of the Qip1 protein. We hypothesize that this modestly truncated Qip1 protein is still functional (Table 1). Taken together, these results indicate that five of the natural isolates identified by the bioinformatic pipelines, strains Bt210, LP-RSA2296, A2-102-5, D17-1, and NRHc5028, lack siRNAs necessary for RNAi silencing.

### Hypermutation requires both RNAi-deficiency and elevated transposon burden

We have previously identified and characterized two clinical isolates (Bt65 and Bt81) with a nonsense mutation in the known RNAi component Znf3, which display a hypermutator phenotype when selected for rapamycin and FK506 resistance due to the increased mobilization of the retrotransposon Cnl1 into *FRR1*, which encodes FKBP12, the shared target of these drugs (43). To determine if the newly identified RNAi-deficient isolates are also hypermutators, fluctuation assays were performed to quantify their mutation rates on medium containing rapamycin and FK506 (R+F) (Fig. 2 and SI Appendix, Dataset S2). None of these isolates showed significantly higher mutation rates on YPD medium containing R+F than the reference strain H99. They also had mutation rates comparable to H99 on YNB medium containing 5-FOA. Only the *msh2*Δ positive control strain, which lacks a core component of the DNA mismatch repair pathway, exhibited a significantly higher mutation rate than H99 on either media (Fig. 2 and SI Appendix, Dataset S3).

**Figure 2.**
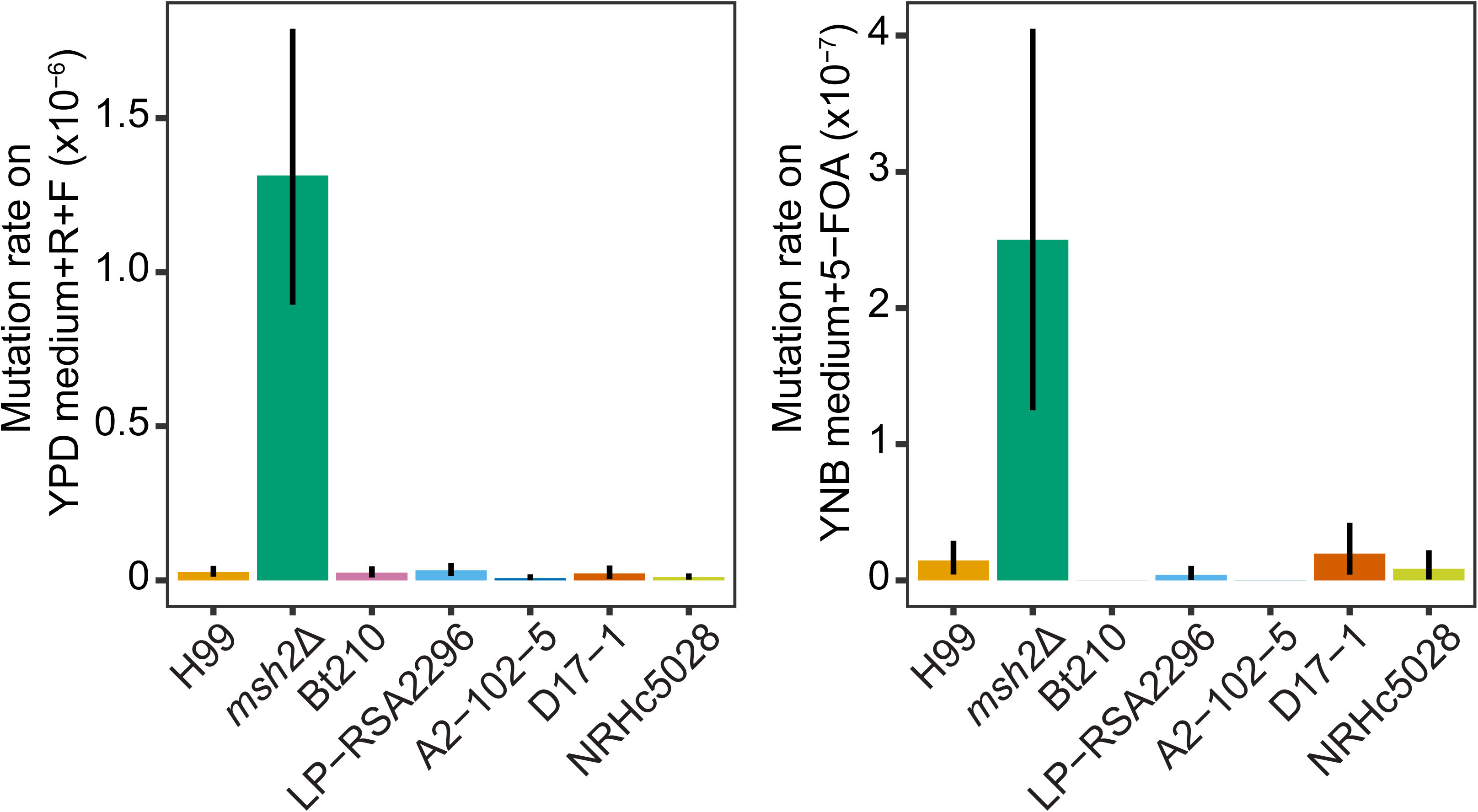
Five novel RNAi-loss isolates do not display a hypermutator phenotype. Mutation rates of five newly identified RNAi-loss isolates when grown on YPD medium containing rapamycin and FK506 (R+F) (left) or YNB medium containing 5-FOA (right), with H99 wildtype and *msh2*Δ strains serving as the negative and positive controls, respectively. The error bars represent 95% confidence intervals.

Given the important role of Cnl1 in causing R+F resistance, we hypothesized that although the five novel isolates had lost functional RNAi, they might not have undergone Cnl1 amplification as observed in Bt65 and Bt81 and, therefore, did not display a hypermutator phenotype. To test this hypothesis, publicly available short-read whole-genome sequence data were analyzed to assess the copy number of full-length Cnl1 (See Materials and Methods) in these isolates. Compared to 150 or 84 copies predicted in Bt65 and Bt81, respectively, the estimated Cnl1 copy number in these isolates ranged from 0 copies (Bt210, A2-102-5, and NRHc5028) to 1 or 3 copies (D17-1 and LP-RSA2296) (Table 2). To validate these short-read based estimations, nanopore long-read sequencing was performed to obtain telomere-to-telomere assemblies for Bt210 and LP-RSA2296. Relative to Bt65 and Bt81, fewer full-length copies of Cnl1 (0 copies in Bt210 and 1 copy in LP-RSA2296) were confirmed by high-quality genome assemblies. These results provide evidence that being RNAi-deficient alone is likely not sufficient for hypermutation, and a relatively high transposon burden is likely also required.

**Table 2.** Full-length Cnl1 copy numbers.

| Isolate | Reference-based assembly | Short-read-based estimation<sup>1</sup> | Hypermutator when selected on YPD+R+F? |
| --- | --- | --- | --- |
| H99 | 0 | 0 | No |
| Bt65 | 105 | 150 | Yes |
| Bt81 | 89 | 84 | Yes |
| Bt210 | 0 | 0 | No |
| LP-RSA2296 | 1 | 3 | No |
| A2-102-5 | NA | 0 | No |
| D17-1 | NA | 1 | No |
| NRHc5028 | NA | 0 | No |

<sup>1</sup>: Whole-genome sequence data from Desjardins et al., 2017 were used for estimation (28).
NA: not applicable; R+F: rapamycin+FK506

Given the observed correlation between Cnl1 burden and hypermutator phenotype, we hypothesized that introduction of the novel RNAi LOF mutations identified here into a strain with a high Cnl1 burden would result in a hypermutator phenotype. In our previous study, restoration of functional RNAi against Cnl1 was successful by repairing the *znf3* nonsense mutation (to result in isolate Bt65+*ZNF3*) (43). Bt65+*ZNF3* contains a Cnl1 burden like that of Bt65 but is not a hypermutator due to its restored RNAi pathway, making it an ideal strain for testing this hypothesis. The *rde4* splice-site mutation from Bt210 and the *znf3* nonsense mutation from LP-RSA2296 (which differs from the original *znf3* nonsense mutation in Bt65, SI Appendix, Fig. S1A) were introduced into the corresponding gene in Bt65+*ZNF3* via CRISPR-mediated allele exchange. One *rde4* transformant and two independent *znf3* transformants were successfully generated (SI Appendix, Fig. S2). In addition to the desired mutation, Sanger sequencing and Plasmidsaurus amplicon sequencing confirmed no additional mutations were found in these two loci. The *rde4* splicing error was also confirmed in Bt65+ZNF3 *rde4* by RT-PCR and Sanger sequencing (SI Appendix, Fig. S2). sRNA-seq further confirmed that like Bt65+*ZNF3 ago1*Δ, the canonical 21-24 nt sRNA peak with 5’-uracil identity was absent in these genetically modified *rde4* and *znf3* mutants, indicating defects in siRNA-mediated RNAi (Fig. 3A and SI Appendix, Fig. S3). To determine whether the introduced mutations confer a hypermutator phenotype in Bt65+*ZNF3*, mutation rates of these re-engineered RNAi-deficient strains were quantified by fluctuation analysis. In alignment with our prediction, when selected for R+F resistance, all three edited strains displayed significantly higher mutation rates than the recipient strain *Bt65+ZNF3* and negative control H99 (Fig. 3B and SI Appendix, Dataset S4).

**Figure 3.**
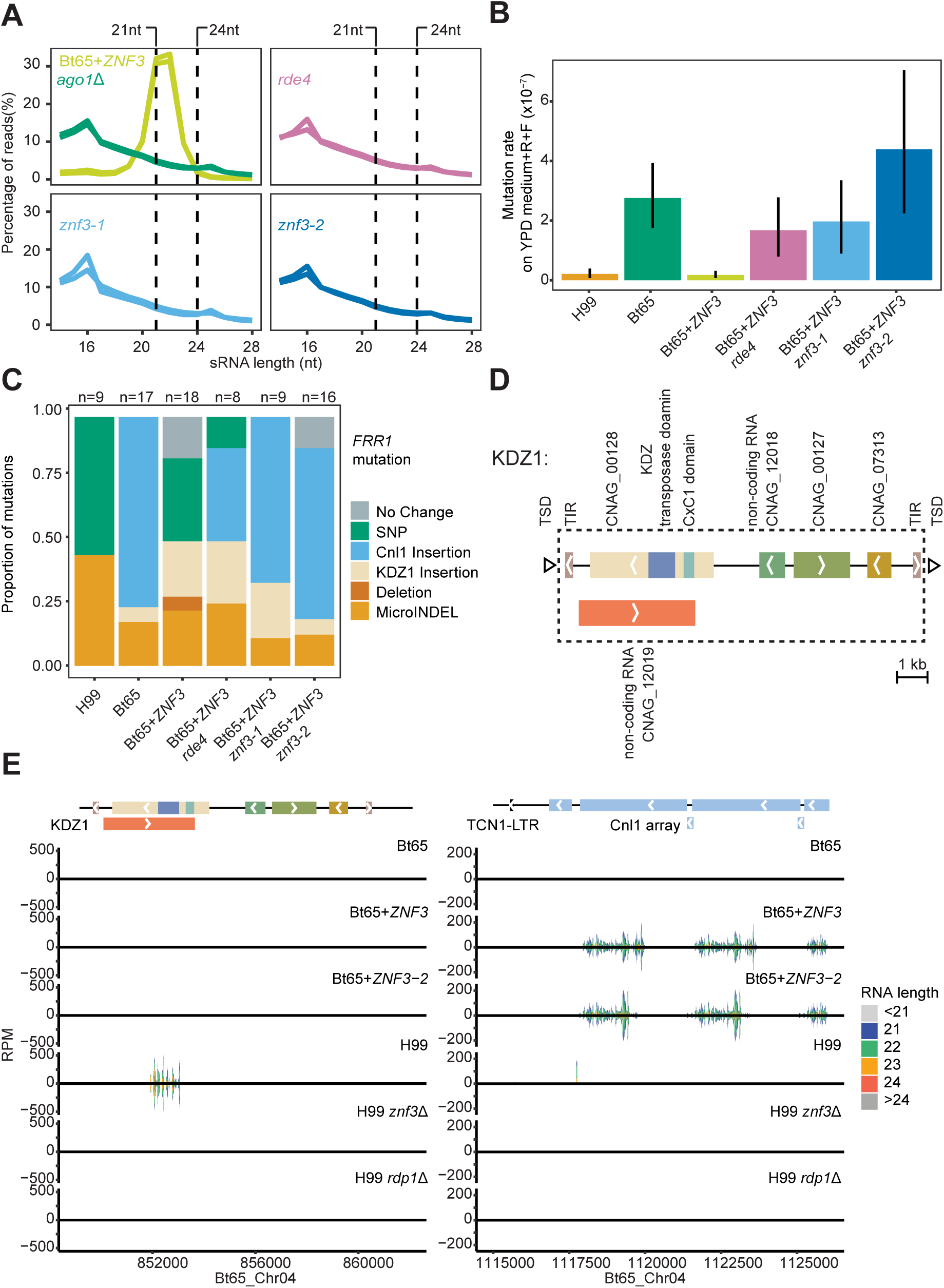
Engineered RNAi loss-of-function mutations result in transposition leading to hypermutation. (A) Size distribution of sRNA reads from each of the respective strains (Bt65+*ZNF3*, Bt65+*ZNF3 ago1*Δ, Bt65+*ZNF3 rde4*, Bt65+*ZNF3 znf3-1*, and *-2*). Two biological replicates of each strain are plotted. All the sRNA reads were mapped to the Bt65 genome. (B) Mutation rates of Bt65+*ZNF3*, Bt65+*ZNF3 rde4* and two independent Bt65+*ZNF3 znf3* allele exchange strains when grown on YPD medium containing rapamycin and FK506, with H99 and Bt65 serving as negative and positive controls, respectively. The error bars represent 95% confidence intervals. (C) Mutational spectra in the *FRR1* gene of independent rapamycin+FK506 resistant colonies from each indicated strain. R+F resistant colonies without mutations detected at the *FRR1* locus are labeled as ‘No change’. (D) Diagram illustrating the structure of a novel gigantic DNA transposon in *C. neoformans*, KDZ1. Genes and non-coding RNAs are shown by solid blocks, with arrows indicating the orientation. Tags starting with “CNAG” correspond to the annotated number in the H99 reference genome. TIR: Terminal Inverted Repeat; TSD: Target Site Duplication (drawn out of scale). (E) Mapping of small RNA reads from Bt65, Bt65+*ZNF3*, Bt65+*ZNF3-2*, H99, H99 *znf3*Δ and H99 *rdp1*Δ aligning to a KDZ1 (left) and a Cnl1 array (right), both of which are located on chromosome 4 of Bt65. Shown on the y-axis are numbers of normalized reads (RPM, Reads Per Million on forward strand (positive value) or reverse strand (negative value)), representing normalized coverage counts at each position. Different lengths of small RNA reads are labeled as indicated by the color.

To determine whether Cnl1 insertion is the main genetic change causing resistance, the FKBP12-encoding gene *FRR1* from spontaneous R+F resistant colonies derived from each strain was PCR amplified and sequenced. Consistent with our previous study, Cnl1 insertions were frequently observed in *FRR1* from the resistant colonies derived from the RNAi-deficient strain Bt65 (13/17, ∼76%), but not the repaired RNAi-proficient strain Bt65+*ZNF3* (0/18, 0%) or H99 (0/9, 0%). In the three re-engineered RNAi-deficient strains, a high portion of R+F resistance was caused by insertion of Cnl1 into *FRR1* (ranging from ∼37.5% to 69%), with the remaining resistant colonies having other types of genetic changes such as single nucleotide polymorphisms (SNPs), microINDELs (insertions or deletions ≤ 50 bp), or deletions (>50 bp deletion) (Fig. 3C and SI Appendix, Dataset S5). Interestingly, relative to the R+F sensitive progenitor, we also observed some resistant isolates from both Bt65+*ZNF3* (3/18, ∼17%) and Bt65+*ZNF3 znf3-2* (2/16, 12.5%) that do not have any mutation (i.e., No change) at the *FRR1* locus including 1,699 bp 5’ and 1,268 bp 3’ sequences, leaving the cause of R+F resistance in these isolates unknown (Fig. 3C and SI Appendix, Dataset S5).

### Movement of a novel DNA transposon occurs in the presence of functional RNAi, giving rise to drug resistance

During the fluctuation assay of the re-engineered RNAi-deficient strains, we noticed that some R+F resistant colonies had *FRR1* alleles containing insertions obviously larger than the expected size of the full-length Cnl1 element (∼3.5 kb) (SI Appendix, Fig. S4 and SI Appendix, Dataset S5). Amplicon sequencing revealed these unusually large insertions were identical (10,944 bp element) and observed in the Bt65-derived strains regardless of their RNAi status. The frequency of resistant colonies attributable to this 10,944 bp insertion in *FRR1* ranged from ∼6% (1/17 in Bt65) to 25% (2/8 in Bt65+*ZNF3 rde4*) (Fig. 3C). Two hallmark signatures of transposon movement, target site duplication (TSD, 8 bp) and terminal inverted repeats (TIR, 236 bp), were detected (Fig. 3D), suggesting the presence of a large uncharacterized mobile element. Although this element was not identified as mobile in H99, BLASTn results suggest the presence of a similar copy (100% query coverage and 98.41% identity) in the H99 genome. Based on the annotation of the H99 reference genome, two non-coding RNAs and three coding genes are included in this region; among these CNAG_00128 is predicted to contain a KDZ (Kyakuja, Dileera, and Zisupton) transposase domain and a putative zinc chelating domain, CxC1, indicative of a KDZ DNA transposon (51) (Fig. 3D). CNAG_12018, and CNAG_12019 are predicted non-coding RNAs, of which CNAG_12019 overlaps with CNAG_00128 in the opposite orientation. The other two genes, CNAG_00127 and CNAG_07313, encode unknown protein products (Fig. 3D). Due to its mobility and canonical KDZ transposon domain structure, we named this novel DNA transposon KDZ1 (the **1**^st^ recognized **KDZ** element) in *C. neoformans*, whose movements have not been previously documented in pathogenic *Cryptococcus* species. In all cases of KDZ1 insertion into *FRR1*, full-length copies were observed (SI Appendix, Dataset S5).

The observation of KDZ1 transpositions in the repaired RNAi-proficient strain Bt65+*ZNF3* suggests that KDZ1 movement might evade functional RNAi machinery. To investigate whether RNAi is involved in silencing KDZ1, we re-analyzed our published sRNA-seq dataset for the RNAi-deficient strain Bt65, two independent RNAi-repaired strains (Bt65+*ZNF3* and Bt65+*ZNF3-2*), as well as H99 and its derived RNAi-deficient mutants H99 *rdp1*Δ and H99 *znf3*Δ (43). The sRNA reads were first mapped against an array of Cnl1 elements located at the 3’ end of the Bt65 Chr4. Abundant 21 to 24 nt sRNA reads targeting Cnl1 in this region were present in the RNAi-repaired strains Bt65+*ZNF3* and Bt65+*ZNF3-2*, while no such peaks were observed in the RNAi-deficient Bt65 or other H99-derived strains lacking a full-length Cnl1 (Fig. 3E). In contrast, no obvious sRNA targeting KDZ1 on the same Bt65 Chr4 was detected in Bt65+*ZNF3* or Bt65+*ZNF3*-2. The only enrichment of sRNA reads was found in H99 around the overlapping region of CNAG_00128 and the anti-sense non-coding RNA CNAG_12019, which also covers the KDZ transposase domain. This enrichment is abolished in H99 *rdp1*Δ and H99 *znf3*Δ, indicating a possible role for these sRNAs in RNAi silencing (Fig. 3E).

Our observation that the movement of a novel KDZ DNA transposon could lead to antifungal resistance in *C. neoformans* prompted us to conduct further detailed investigation into the diversity of KDZ transposons. Specifically, we first conducted a Hidden Markov Model (HMM) search using the KDZ transposase domain (PF18758) against the genomes of the reference strain H99 and the hypermutator strain Bt65 and identified five and six homologous sequences with high-confidence E-values, respectively (SI Appendix, Table S2). We then conducted phylogenetic analysis of these KDZ domain-containing proteins, together with those from the known KDZ transposons from the model mushroom *Coprinopsis cinerea* that belong to the clades of Kyakuja, Dileera, and Zisupton (Fig. 4A). Our results suggest that all the KDZ domain-containing proteins in H99 and Bt65 belong to the Dileera clade with strong bootstrap support. The only exception was CNAG_07523, whose placement in between the Dileera and Kyakuja clades still needed to be further resolved (Fig. 4A). It should be noted that for the vast majority of the KDZ domain-containing proteins in H99 and Bt65, we did not detect in their vicinity any of the other features that are characteristic of a complete KDZ transposon, such as the CxCn domain, as well as the TIRs and/or TSDs flanking the element, suggesting they may not represent active mobile elements. Nevertheless, we did identify two KDZ-containing proteins in the Bt65 genome (termed KDZ2 and KDZ3, which are absent in the H99 genome) that are associated with the CxC1 domain, residing in large DNA segments (10,053 bp for KDZ2 and 9, 218 bp for KDZ3) with TSDs and TIRs on both ends (Fig. 4B). Sequence analyses suggest that the KDZ proteins in KDZ2 and KDZ3 are significantly different from each other, although they are evolutionarily more closely related to each other than they are to KDZ1 (Fig. 4A and 4B). Indeed, we found one KDZ2 insertion in *FRR1* from a Bt65-derived F1 progeny (SI Appendix, Fig. S5). Thus, it is reasonable to propose that there are three different types of full-length KDZ transposons in Bt65, all of which are mobile and can induce genetic changes through their transposition.

**Figure 4.**
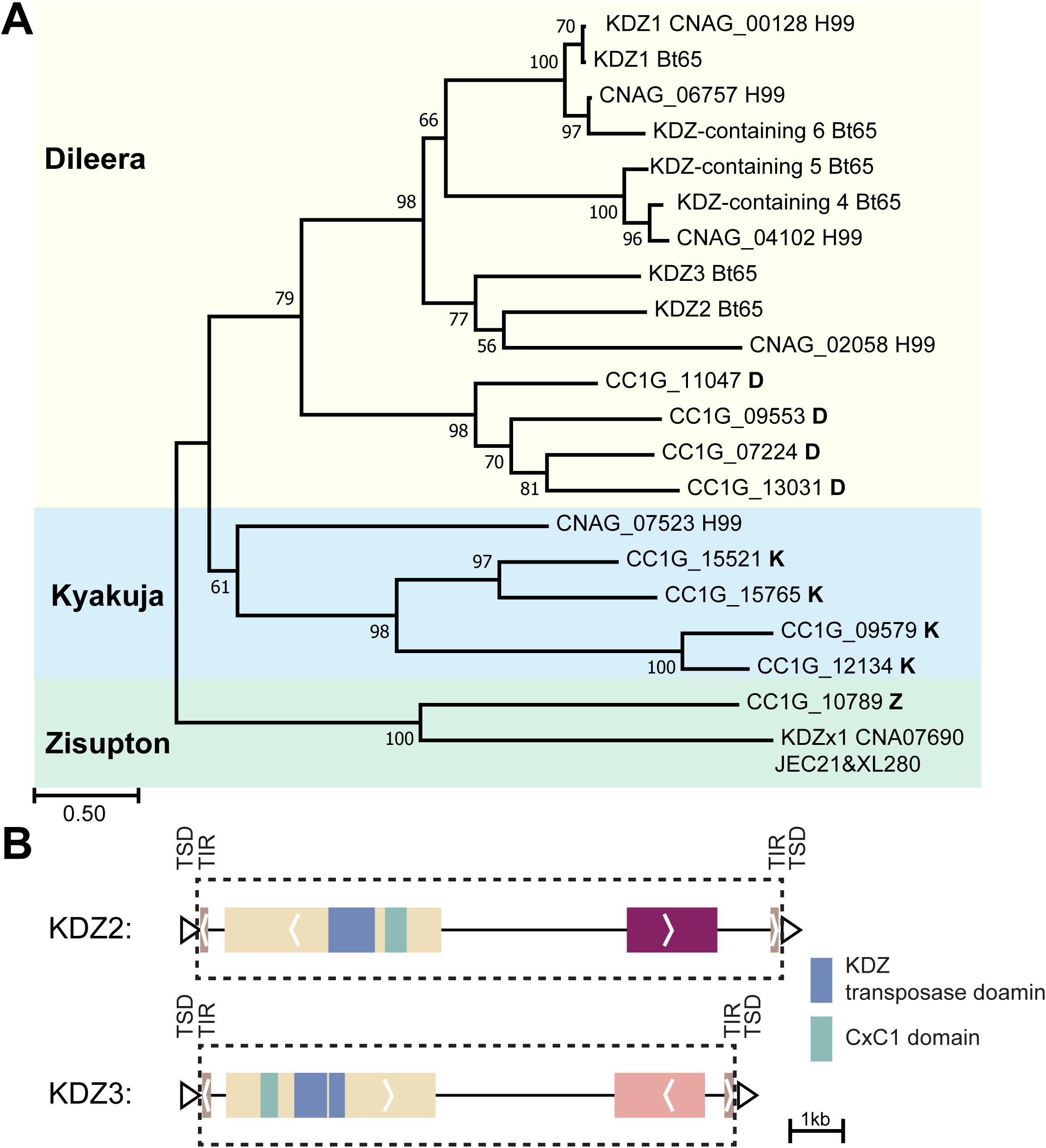
Evolutionary diversity of the giant KDZ DNA transposons in *Cryptococcus*. (A) Maximum-likelihood phylogeny of KDZ domain-containing proteins from Bt65 and H99, as well as representative KDZ domain-containing proteins that have been characterized from *Coprinopsis cinerea* (CC1G**_**). The tree was evaluated with 1000 bootstrap replicates. Shades of different colors highlight clades containing the *Coprinopsis cinerea* Kyakuja (K), Dileera (D), Zisupton (Z) proteins, respectively. (B) Diagram illustrating the structure of two novel gigantic DNA transposons in *C. neoformans*, KDZ2 and KDZ3. Genes are shown by solid blocks, with arrows inside indicating the orientation. TIR: Terminal Inverted Repeat; TSD: Target Site Duplication (drawn out of scale).

### Transposon accumulation during experimental evolution leads to hypermutation in *Cryptococcus*

We identified a strong correlation between the hypermutator phenotype and TE burden (e.g., Cnl1 burden) in the absence of functional RNAi. We next tested if a transition from non-hypermutator to hypermutator could occur following *in vitro* passage, a process known to lead to accumulation of TEs in *C. deneoformans*, a sister species of *C. neoformans* (52). Specifically, we selected two natural isolates that are RNAi-deficient and have low TE burdens in their genomes, Bt210 and LP-RSA2296, and conducted serial passages of the two strains, together with the *C. neoformans* reference strain H99 as control, on YPD solid medium for 2 months at 30°C and 37°C (transferred as patches every two days). Fluctuation assays were then conducted to estimate mutation rates on YPD+R+F solid medium for strains after 28 passages (mutation accumulation 28, i.e., MA28) and 56 passages (i.e., MA56). None of the passaged strains showed increased mutation rates on YPD+R+F solid medium compared to the H99 reference strain or the B210 and LP-RSA2296 progenitor strains (SI Appendix, Fig. S6A; SI Appendix, Dataset S6), even though an initial pilot assay of these strains by a spread plate method suggested that Bt210-MA28 at 30°C, as well as LP-RSA2296-MA28 at both 30°C and 37°C, might have had increased mutation rates compared to the control strains (SI Appendix, Fig. S7). The mutational spectra of R+F resistant colonies from Bt210, LP-RSA2296, as well as their passaged strains were further characterized. We did not observe any TE insertions in *FRR1*, and instead SNPs, deletions, and MicroINDELs were observed (SI Appendix, Fig. S6B and SI Appendix, Dataset S7). Additionally, we estimated the copy numbers of Cnl1 and KDZ1 in the passaged strains using Illumina whole-genome sequencing reads and found no detectable accumulation of Cnl1 or KDZ1 in the passaged strains relative to their progenitor strains (SI Appendix, Table S3). These results suggest that the RNAi-loss isolates have not undergone transposon amplification following a short period of *in vitro* passage and, consequently, did not display a hypermutator phenotype.

We further tested this hypothesis in *Cryptococcus deneoformans*, a sister species of *C. neoformans*. In this case, we utilized a strain, 37-02, which was derived from the laboratory strain XL280α by serial *in vitro* passage (52). Previous studies have shown that compared to the parental strain XL280α, strain 37-02 has undergone expansion of the Cnl1 retrotransposon (with ∼40 additional copies) as well as other transposons in its genome (52). We also analyzed XL280α *rdp1*Δ and generated 37-02 *rdp1*Δ deletion strains to study the effects of RNAi on the mutation rates in these two genetic backgrounds and confirmed that they are indeed RNAi-deficient based on sRNA-seq analysis (Fig. 5A and SI Appendix, Fig. S8). Using fluctuation assays for R+F resistance, the passaged strain 37-02 had significantly higher mutation rates compared to its progenitor strain XL280α, suggesting that the expansion of transposons in the genome likely transformed strain 37-02 into a hypermutator following *in vitro* passage (Fig. 5B, SI Appendix, Dataset S8). Strain XL280α *rdp1*Δ had a mutation rate that was significantly higher than XL280α and comparable to the passage strain 37-02, indicating that inactivation of the RNAi machinery in this genetic background has a similar effect on mutation rates compared to transposon copy number expansion (Fig. 5B). The two independent *rdp1*Δ deletion strains in the 37-02 strain background exhibited mutation rates considerably higher than the XL280α *rdp1*Δ strain, and significantly higher than that of the 37-02 passage strain, providing evidence that the impacts of RNAi loss and TE accumulation on mutation rate are additive (Fig. 5B).

**Figure 5.**
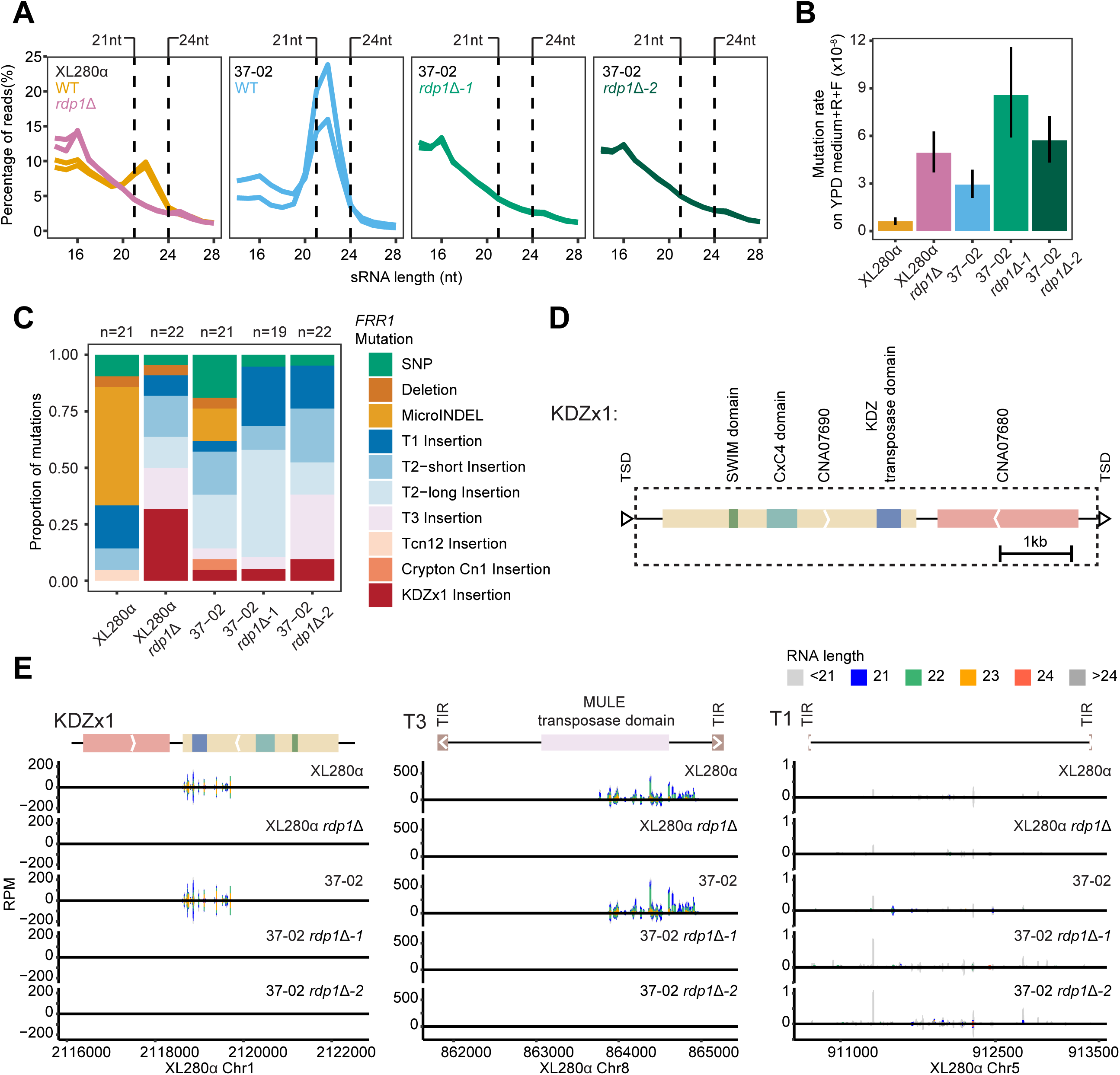
Dynamic outcomes of short-term experimental evolution. (A) Size distribution of sRNA reads from each indicated strain. sRNA reads from two biologically independent replicates are plotted. All the sRNA reads were mapped to the XL280α genome. (B) Mutation rates of XL280α, XL280α *rdp1*Δ, 37-02, 37-02 *rdp1*Δ*-1*, and *-2* when grown on YPD medium containing R+F are shown. (C) Mutational spectra in the *FRR1* gene of independent R+F resistant colonies from each indicated *C. deneoformans* strain. (D) Diagram illustrating the structure of a novel DNA transposon in *C. deneoformans*, KDZx1. Solid colored blocks represent open reading frames, with arrows inside indicating the orientation. CNA07680 and CNA07690 correspond to two proteins in the *C. deneoformans* JEC21 reference genome. TIR: Terminal Inverted Repeat; TSD: Target Site Duplication (drawn out of scale). (E) Mapping of small RNA reads from XL280α, XL280α *rdp1*Δ, 37-02, 37-02 *rdp1*Δ*-1*, and *-2* to a KDZx1 (left), a DNA transposon T3 (middle, with MULE transposase domain and TIRs), and a DNA transposon T1 (right, only known with TIRs), which are located on different chromosomes of XL280α. Shown on the y-axis are normalized reads (RPM, Reads Per Million on forward strand (positive value) or reverse strand (negative value)), representing normalized coverage counts at each position. Different lengths of small RNA reads are labelled as indicated by the color.

We next analyzed the mutation spectra of the R+F resistant colonies from all the tested strains by PCR and sequence analysis of the *FRR1* gene. With the exception of the wild-type progenitor strain XL280α, where the majority of mutations in the *FRR1* gene were due to changes other than transposon insertions (e.g., SNPs, deletions, and MicroINDELs), insertions of various transposable elements into the *FRR1* gene were responsible for the majority of R+F resistance in all of the other four derived and genetically modified strains tested, including 37-02 (Fig. 5C). In total, seven different types of transposons were detected via insertions into the *FRR1* gene. Interestingly, one of the mobile elements is a 6,397 bp novel element capped by 8 bp target site duplications on both ends; insertion of this element occurred in all strains tested except XL280α. Further analyses showed that this element contains two open reading frames based on the annotation of the *C. deneoformans* type strain JEC21 (CNA07680 and CNA07690). While CNA07680 encodes an unknown protein, the protein encoded by CNA07690 contains a KDZ transposase domain, a CxC4 domain, and a SWIM (SWI2/SNF2 and MuDR) domain, which fits the canonical domain structure of the Zisupton transposon from the KDZ DNA transposon class (8, 12) (Fig. 5D). We therefore named this novel DNA transposon KDZx1 (**KDZ** in **X**L280 #**1**) in *C. deneoformans* (Fig. 4A, Fig. 5C and D). Additional analysis showed that this element is also present in the *C. deneoformans* strain JEC21, although it should be noted that in both cases we did not detect terminal inverted repeats that typically are located at the two ends of KDZ transposons (Fig. 5C, D and SI Appendix, Dataset S9). Analysis of complete genome assemblies of strains XL280α and 37-02 showed that KDZx1 had not undergone copy number expansion during *in vitro* passage (SI Appendix, Table S4).

Surprisingly, we did not detect any Cnl1 insertions in *FRR1* among the 105 R+F resistant colonies analyzed in the *C. deneoformans* strains (Fig. 5C and SI Appendix, Dataset S9). We hypothesize that different transposons might be regulated by diverse mechanisms (e.g., RNAi-dependent vs. RNAi-independent). To test this hypothesis, we counted the coverage of sRNA reads in multiple transposons and found that KDZx1 and the DNA transposon T3 are enriched with 21 to 24 nt sRNA reads near their transposase domain, while no such enrichment was observed for the DNA transposon T1 (Fig. 5E). Consistent with this finding, there is significant evidence to suggest that the transposition of KDZx1 (Fisher’s Exact Test, p-value= 0.04671) and T3 (Fisher’s Exact Test, p-value= 0.02531) is different between RNAi-proficient and RNAi-deficient backgrounds (Fig. 5C and SI Appendix, Dataset S9). However, there is no such correlation for the movement of T1 (Fisher’s Exact Test, p-value= 0.5819) (Fig. 5C and SI Appendix, Dataset S9). Taken together, our results show that both RNAi loss and/or transposon accumulation can lead to hypermutation in *Cryptococcus* via uncontrolled transposition.

### Hypermutation caused by RNAi loss and transposition can occur following sexual reproduction

We have identified a new class of mobile elements in *C. neoformans*, the KDZ transposons, whose transposition can give rise to antifungal resistance. It is known that transposons are activated during meiosis, and genetic variation resulted from sexual reproduction and meiotic recombination could have contributed to the observed variation in KDZ and Cnl1 distribution and activity across isolates (47, 53). Thus, we sought to investigate how sexual reproduction could influence: 1) the activity and regulation of transposons, and 2) the mutation rates of RNAi-deficient progeny inheriting a transposon burden. To this end, we crossed two natural isolates, recovered meiotic progeny, and studied the mutation rates, and their relationship to transposon (e.g., KDZ and Cnl1 transposon) movements and regulation. Specially, we crossed strain Bt210 (*ZNF3 rde4*) with strain Bt65 (*znf3 RDE4*). Bt210 is RNAi-deficient due to a mutation in the *RDE4* gene and a non-hypermutator likely due to a lack of sufficient transposon burden in the genome. Bt65 is an RNAi-deficient hypermutator because of a nonsense mutation in a key RNAi gene, *ZNF3*, and with an elevated load of transposable elements in its genome (e.g., KDZ and Cnl1 elements). A total of 57 meiotic progeny were recovered, and segregation analyses of *ZNF3*, *RDE4*, and the mating type locus confirmed that this progeny set represents a population of random meiotic products (SI Appendix, Dataset S10). Whole-genome sequencing also confirmed meiotic recombination on a genome-wide scale (SI Appendix, Fig. S9).

Among the tested F1 progeny (all 11 progeny with *rde4 ZNF3* and 2 progeny with *rde4 znf3*), three exhibited a hypermutator phenotype on YPD medium containing R+F. Two of these hypermutators (P12 and P25) were *ZNF3 rde4* (same genotype as the Bt210 parent), while the other one (P57) was *znf3 rde4*, a recombinant genotype between the Bt210 and Bt65 parents. While all three F1 progeny had mutation rates that were significantly higher than the non-hypermutator parent Bt210, when compared to that of the hypermutator parent Bt65, the mutation rate of P57 was relatively lower while those of P12 and P25 were comparable to each other (albeit lower than Bt65 as well) (Fig. 6A and SI Appendix, Dataset 11). sRNA sequencing confirmed that the three hypermutator F1 progeny (P12, P25, and P57) are RNAi-deficient, lacking the 21 to 24 nt sRNA species and the 5’-uracil enrichment in sRNA reads (Fig. 6B and SI Appendix, Fig. S10).

**Figure 6.**
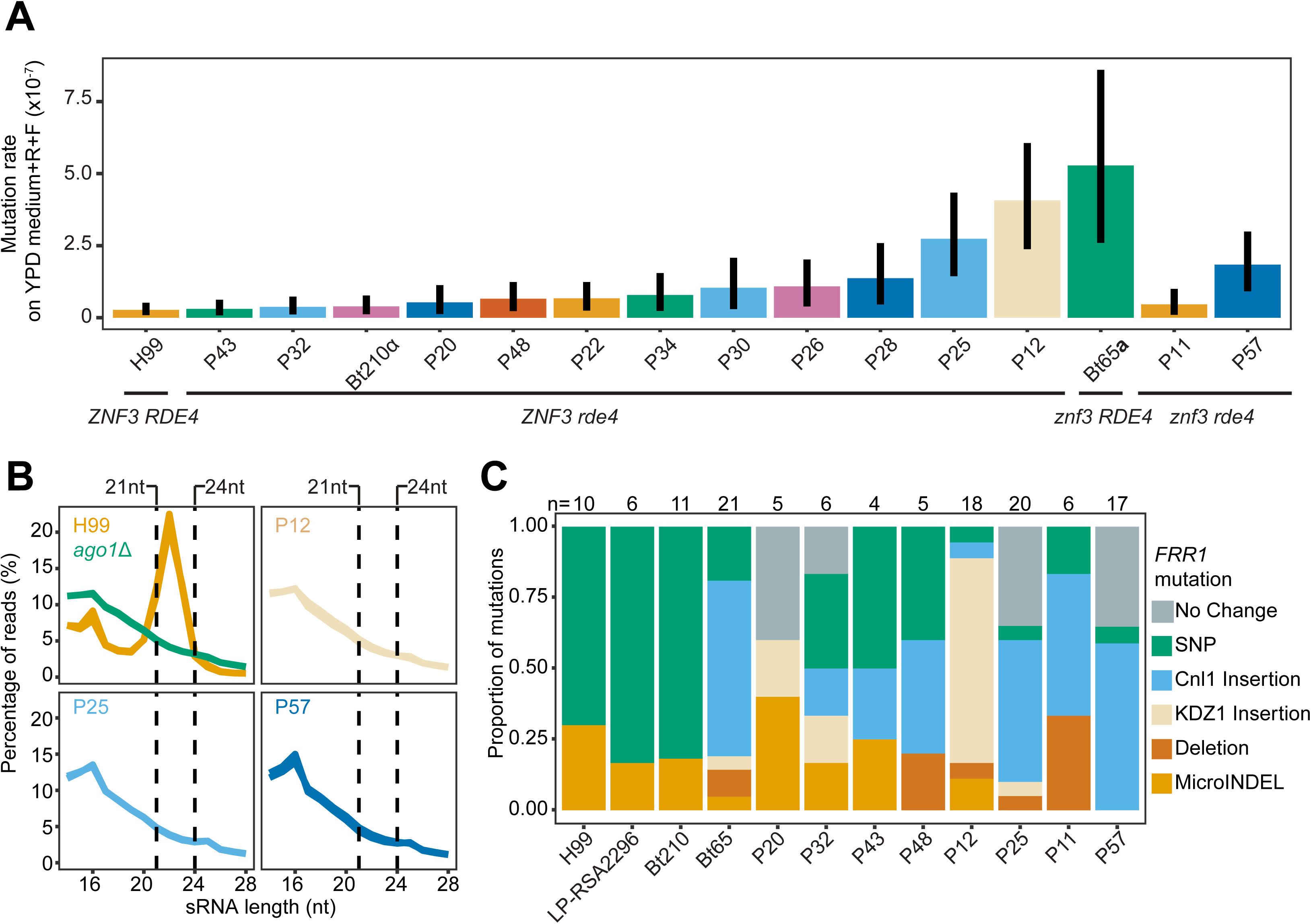
Transposition causes hypermutation after sexual reproduction. (A) Mutation rates of 13 F1 progeny dissected from the cross between natural isolates Bt65**a** and Bt210α, with the two parents (Bt65**a** and Bt210α) and *C. neoformans* reference strain H99 serving as controls, when grown on YPD medium containing rapamycin + FK506. The error bars represent 95% confidence intervals. Genotypes of *RDE4* and *ZNF3* are shown for each strain. (B) Size distribution of sRNA reads from each indicated strain with two independent biological replicates for each strain. All the sRNA reads were mapped to the Bt65 genome. (C) Mutational spectra in *FRR1* in independent rapamycin + FK506 resistant colonies from each indicated strain.

The mutation spectra at the *FRR1* locus in R+F resistant colonies derived from control, parental, and both hypermutator and some non-hypermutator progeny were assessed (Fig. 6C). Insertion of the KDZ1 and Cnl1 transposons at the *FRR1* locus contributed to a considerable proportion of R+F resistant isolates in the hypermutator parent Bt65, as well as in the F1 progeny regardless of whether they were hypermutators or not. This is in stark contrast to the non-hypermutator parent Bt210 as well as H99 and LP-RSA2296, where the vast majority of the R+F resistant colonies had SNPs in the *FRR1* gene, and no transposon insertions were detected. Interestingly, while in Bt65, the majority of transposon insertions were attributable to the Cnl1 element with few KDZ1 insertions being observed (∼5%, 1 of 21 independent colonies). A relatively elevated number of KDZ1 insertions were observed in three F1 progeny (P20, P32, and P12), of which P12 is a hypermutator with 13 of 18 (∼72%) independent resistant colonies derived from this progeny contained KDZ1 insertions in *FRR1* (Fig. 6C and SI Appendix, Dataset 12). Importantly, in the two other hypermutator progeny (P25 and P57), the majority of R+F resistant colonies (50%, 10/20 in P25 and ∼59% 10/17 in P57) were attributable to Cnl1 insertions into the *FRR1* gene. Additionally, there were resistant colonies derived from P25 (35%, 7/20) and P57 (∼35%, 6/17) in which we could not detect any mutation at the *FRR1* locus (including 1,699 bp 5’ and 1,268 bp 3’ sequences). To test the correlation between the hypermutator phenotype and the inherited transposon burden in the F1 progeny, we estimated the copy numbers of Cnl1 and KDZ1 by mapping the Illumina short-reads. We found that collectively the three hypermutator F1 progeny inherited significantly more Cnl1 elements than the remaining non-hypermutator progeny (Mann-Whitney U Test, p-value= 0.03571) (SI Appendix, Table S5). Moreover, progeny P12, which had the highest frequency of KDZ1 insertion among resistant colonies, also contained the highest abundance of KDZ1 (7 copies), which is higher than any other tested progeny as well as the two parents (SI Appendix, Table S5). Thus, our results demonstrate that sexual reproduction has the potential to 1) liberate the KDZ mobile elements and 2) cause the inheritance of a high Cnl1 element burden along with a loss of RNAi mutation, resulting in hypermutation.

To determine whether hypermutation can result in resistance to clinically relevant antifungal drugs, mutation rates on medium containing 5-fluorocytosine (5-FC), a systematic antifungal that is one of the first-line treatments for cryptococcal infection, were measured. Fluctuation analysis suggested that none of the three F1 progeny as well as Bt210 and Bt65 display a hypermutator phenotype when selected for 5-FC resistance (SI Appendix, Fig. S11). We further analyzed the 5-FC resistant colonies isolated from P12, P25, P57, and the parents Bt210 and Bt65 by PCR amplifying three genes in which mutations are known to confer 5-FC resistance (*FUR1*, *UXS1* and *FCY2*) (54). Sequencing results revealed that KDZ1 insertions underlie resistance in four 5-FC resistant colonies derived from P12. Two insertions were observed in *FUR1*, and two were observed in *UXS1*. Additionally, we detected a KDZ1 insertion in *UXS1* in one of the resistant colonies derived from Bt65 (SI Appendix, Fig. S11 and SI Appendix, Dataset 13). These results demonstrate that KDZ1 can mediate resistance to a clinically relevant antifungal drug.

## Discussion

Two bioinformatic pipelines were developed to identify mutations in required RNAi components using short-read whole-genome sequence data for 387 *C. neoformans* natural isolates (28). In addition to two previously identified RNAi-loss hypermutators, Bt65 and Bt81, these pipelines successfully identified five novel RNAi-deficient isolates (Bt210, LP-RSA2296, A2-102-5, D17, and NRHc5028), as evidenced by sRNA-seq analysis. In contrast to Bt65 and Bt81 (from the Sub-Saharan African VNBII lineage), all five newly identified RNAi-loss isolates are from the global VNI lineage; among these, clinical isolates Bt210 and LP-RSA2296 belong to two different clades, while environmental isolates A2-102-5, D17-1, and clinical isolate NRHc5028, are all from a single clade. The identification of five additional non-VNB RNAi-deficient isolates in *C. neoformans* suggests that the loss of RNAi within this species may be more widespread than previously appreciated. To date, RNAi loss has been confirmed in seven natural isolates, which account for ∼1.8% of the 387 isolates in the Strain Diversity Collection (28, 43). Given our limited understanding of the RNAi components in *C. neoformans* and the nature of our pipeline (from genotype to phenotype and only focused on LOF mutations), the full extent of RNAi loss in this species may still be underestimated. Interestingly, a recent study revealed the presence of an intact and functional RNAi pathway in the human fungal pathogen *C. albicans*, which was previously thought to be RNAi-deficient based on a single reference strain (SC5314) (25). Our results, along with this recent finding from *C. albicans,* provide robust examples of the importance of variation not only between species, but also within a species and highlight the limitations of making conclusions based on analysis of a single reference strain.

All five newly identified RNAi-deficient isolates failed to display hypermutator phenotypes when selected for R+F or 5-FOA resistance. Compared to the hypermutator isolates Bt65 and Bt81, which have approximately 100 copies of the non-LTR retrotransposon Cnl1, the five newly identified RNAi-deficient isolates contain significantly fewer copies of Cnl1 (0 to 4 copies). This association between the hypermutator phenotype and copy number of the Cnl1 retrotransposon is in line with our hypothesis that it is the combination of RNAi loss and a high transposon burden that causes hypermutation (43). This hypothesis is also supported by the transition from non-hypermutator to hypermutator observed in our CRISPR re-engineered allele exchange strains. These findings elucidate two distinct evolutionary trajectories following RNAi loss: one pathway leading to accumulation of transposons and resulting in a hypermutator phenotype, and the other where the organism maintains or loses transposons and persists as a non-hypermutator (SI Appendix, Fig. S12). Consistent with our model, the overall repeat content is similar between the RNAi-deficient isolates Bt210 and LP-RSA2296, and the RNAi-proficient reference strain H99 (SI Appendix, Dataset 14). Additionally, the loss of RNAi in the *C. deuterogattii* type strain R265 is associated with a reduction of full-length retroelements at the centromeres (24, 44). Although we did not observe significantly increased mutation rates or transposon accumulation following passage of RNAi-loss strains Bt210 and LP-RSA2296, possibly attributable to the relatively short passage time and low initial transposon numbers (0 copies Cnl1 in Bt210 and 1 copy of Cnl1 in LP-RSA2296), it remains possible that the transposon burden could increase over time without RNAi and eventually lead to hypermutation.

Our results in *C. deneoformans* indicate that even with functional RNAi, transposon accumulation can lead to hypermutation, as seen by the increased mutation rate of the passaged strain 37-02 compared to its progenitor XL280α (52). Additionally, removing RNAi in 37-02 further increased the mutation rate, highlighting an additive effect between RNAi loss and transposon accumulation. This suggests an alternate evolutionary path to become a hypermutator in which transposons first accumulate followed by RNAi loss (SI Appendix, Fig. S12). Interestingly, drug-resistant colonies with various types of transposon insertions were found in both RNAi-proficient XL280α and 37-02, as well as in their RNAi-deficient deletion mutants. This observation, combined with distinct sRNA read enrichment, suggests the presence of RNAi-independent mechanisms controlling transposable elements and highlights a substantial transposon reservoir in *C. deneoformans*. Furthermore, unlike Bt65 and Bt81 (43), after screening more than 100 independent R+F resistant colonies from *C. deneoformans*, we did not find any Cnl1 insertions in the *FRR1* gene. These results suggest a distinct insertion preference of Cnl1 between *C. neoformans* and *C. deneoformans* and might also be due to their difference in Cnl1 copy number. Similarly, a previous study found no evidence of Cnl1 moving into non-telomeric regions when selected for antifungal drug resistance in *C. deneoformans* (52).

During our efforts to characterize mutational spectra, we surprisingly identified a novel class of large KDZ DNA transposons in *Cryptococcus*, KDZ1, KDZ2, KDZ3 in *C. neoformans* and KDZx1 in *C. deneoformans*, whose movements had not been previously documented. We found evidence indicating KDZ1, KDZ2, as well as KDZx1 can mobilize into genes encoding drug targets, thereby giving rise to antifungal drug resistance. Given these observations, it is reasonable to hypothesize that KDZ3 may similarly insert into genes encoding antifungal drug targets. The KDZ DNA transposon class, which consists of three different clades: **K**yakuja, **D**ileera and **Z**isupton, was first identified in the mushrooms *Coprinopsis* and *Laccaria* and shown to be associated with the expansion of TET/JBP dioxygenases in basidiomycetes (51). Our discovery that diverse KDZ transposons are present in *Cryptococcus*, as well as their transposition leading to drug resistance and elevated mutation rates, echoes the growing appreciation of the diversity of transposable elements, and the effects of their movements on the host phenotype and genome integrity and stability in a variety of organisms (42, 55, 56). The average size of our identified KDZ transposons is ∼9.1 kb (ranging from ∼6.4 kb for KDZx1 to ∼10.9 kb for KDZ1), which is larger than most known transposons in *Cryptococcus*, such as the ∼3.5 kb full-length Cnl1. In addition to the coding sequence containing the KDZ transposase domain along with its N-terminal zinc chelating domain (CxCn domain), these transposons also harbor extra open reading frames encoding proteins without predicted functional domains. Other recently characterized giant mobile elements (e.g., *Starship*) are known to carry cargo sequences contributing to genomic variability within various fungal species (57–59), and these additional sequences may function in a similar fashion.

Interestingly, we also found additional annotated proteins in *Cryptococcus* genomes that contain the KDZ transposase domain, although they lack other features of KDZ transposons, such as the CxCn domain, and we did not detect sequences indicative of the flanking TIRs and TSDs in their vicinity. In a recent study, one such protein, CNAG_04102, was implicated to be associated with virulence variation in *C. neoformans* based on a genome-wide association study (60). These KDZ-transposase-containing proteins could represent relics of inactivated KDZ transposons, or they could be unique types of KDZ transposons that are yet to be characterized. Further studies are required to gain a better understanding of the characteristics and biology of this novel class of KDZ mobile elements.

In the *C. neoformans* reference strain H99, a non-coding RNA (CNAG_12019) was also annotated inside the KDZ1 transposon, and it is on the anti-sense strand of the region corresponding to the KDZ transposase domain. This suggests a possibility where double-stranded RNA is formed to target the KDZ transposase and initiate RNAi silencing of KDZ1. However, our observations of distinct activities between Cnl1 and KDZ1 in the RNAi-repaired strain, Bt65+*ZNF3*, suggest a distinct regulatory mechanism for KDZ1. Thus, while RNAi plays an important role in regulating Cnl1 (43), a different or additional mechanism might be involved in regulating KDZ1. Similarly, studies in *C. deneoformans* showed that while both KDZx1 and T3 DNA transposons (50) appear to be regulated by RNAi, the T1 DNA transposon remains active despite functional RNAi. These findings highlight the complexity of the regulation of transposable elements, as well as the consequent transposon-mediated antifungal drug resistance, and emphasize the need for further investigation into the regulatory mechanisms controlling various types of transposons in *Cryptococcus* and other species.

Three F1 hypermutator progeny inheriting the *rde4* splice mutation were identified from a genetic cross between Bt210 (*rde4*, RNAi-loss with limited transposons) and Bt65 (*RDE4*, RNAi-loss with enriched transposons). In addition to the Cnl1 insertions that predominated in two F1 progeny (P25 and P57), we found that KDZ1 transposition underlies most of the R+F resistance observed in one of the progeny (P12). Movement of Cnl1 and KDZ1 was also observed in several non-hypermutator F1 progeny, indicating the active status of these transposons. These findings suggest that hypermutation can evolve in two steps: an initial loss of RNAi by mutation followed by sexual reproduction with an isolate with a high transposon burden. F1 progeny from such a cross could display a hypermutator phenotype due to the combined effects of RNAi loss and inheritance of a transposon burden.

Taken together, our results demonstrate that rampant insertion of transposons, including the novel KDZ and Cnl1 mobile elements, conferred a hypermutator phenotype in the RNAi-deficient progeny produced through sexual reproduction, indicating that RNAi loss provides a pathway to hypermutation. The unique transposition profiles of the KDZ elements among these RNAi-deficient progeny suggest the presence of multifaceted and diverse regulatory mechanisms against different transposable elements, which could result from evolutionary pressure to suppress the propagation of selfish mobile elements and defend host genome integrity. Further research on the transgressive KDZ1 movement following sexual reproduction will illuminate the regulatory mechanism(s) controlling KDZ1. Notably, only a limited number of genetic loci and insertion sizes can be captured by the fluctuation analysis and PCR-based genotyping strategy applied in this study. Due to these experimental limitations, the activity and the role of the gigantic KDZ transposons within the genome are likely underappreciated.

Collectively, our findings revealed relatively frequent RNAi loss in *C. neoformans* natural isolates, and many of these may lie on a pathway that could lead to hypermutator evolution mediated by various types of transposons (SI Appendix, Fig. S12). With the discovery of KDZ transposons in *Cryptococcus*, our study sheds light on a darker side of the fungal genome, which could play a significant role in adaptation within fungal populations.

## Materials and Methods

### Pipeline constructions

To identify potential LOF mutations in the Strain Diversity Collection, two independent pipelines (NeoStop and GenesComp) were constructed; details are provided in the SI Appendix.

### Small RNA isolation, sequencing, and analysis

Small RNA isolation and analysis were performed as described previously (43, 61); details are provided in the SI Appendix.

### Fluctuation assay

Fluctuation assays were performed as described previously (43); details are provided in the SI Appendix. Mutation rates and 95% confidence intervals for each strain were calculated using the FluCalc program (https://flucalc.ase.tufts.edu/) (62). As a strict standard, no overlap between respective 95% confidence intervals indicates a significant difference (63).

### Transposon copy number estimates via relative read depth analysis

To estimate transposon copy number from Illumina whole-genome short-read data we carried out the following analysis steps for each strain of interest: 1) Illumina short reads were mapped to the genome of the reference strain H99 using minimap2 (64), and the median genome-wide read depth was estimated using the tool Mosdepth (65); 2) The same reads were mapped to a “transposon genome” consisting of the concatenated sequences of KDZ1, KDZ2, KDZ3, and Cnl1; 3) For the transposon mapped reads, the cumulative read depth distribution output of Mosdepth was used to estimate the read-depth at which a given query sequence had a coverage of at least 0.95; 4) The read depth at coverage 0.95 (step 3) was divided by the genome-wide median read depth (step 2) to give a normalized read-depth estimate.

To provide a benchmark of the performance of our relative read depth estimates of transposon copy number, we used linear regression to fit copy number estimated via BLAT from whole genome assemblies to the read-depth estimated copy number (SI Appendix, Fig. S13). We find strong agreement between the two estimators of copy number, with R^2^ values of the linear models between 0.95 and 0.98 depending on whether estimates for Cnl1 copy number in Bt65 are included or excluded.

## Data Availability

The *de novo* assemblies of Bt210 and LP-RSA2296 have been deposited with accession number PRJNA1138746 in the NCBI Bioproject database. The raw sequence reads for Nanopore sequencing, sRNA-seq and Illumina whole-genome sequencing have also been deposited under the same BioProject accession number. sRNA data presented in Fig. 3E were collected previously and accessed through NCBI Bioproject accession PRJNA749953. The novel KDZ transposons identified in this work have been submitted to GenBank and are available as GenBank number PQ181658 (KDZ1), PQ181659 (KDZx1), PQ181660 (KDZ2), and PQ181661 (KDZ3).

## Acknowledgments

We thank Anna Floyd-Averette for constant support, and all the members of the Heitman Lab for constructive suggestions. We also thank Dr. Asiya Gusa and Anna Mackey (Duke University) for generously sharing the passaged strain 37-02 and XL280α *rdp1*Δ and for discussions and communication of unpublished results on their independent discovery of the KDZ transposons, Dr. Ruiyun Zeng (North Carolina State University) for assistance in data visualization, and Dr. Tim A. Dahlmann and Dr. Minou Nowrousian (Ruhr-Universität Bochum) for suggestions on sRNA data analysis. This study was supported by NIH/NIAID R01 grants AI039115-27, AI050113-20, and AI133654-07. J. Heitman is co-director and fellow of the CIFAR program Fungal Kingdom: Threats & Opportunities. We thank the Madhani laboratory and NIH grant R01 AI100272 for the KN99α *msh2*Δ and *frr1*Δ deletion strains.

## Author Contributions

J. Huang., S.J.P., V.Y., and J. Heitman designed research; J. Huang., C.J.L., S.J.P., Z.X., and S.S. performed research; J. Huang., P.M.M., C.J.L., and V.Y. analyzed data; P.M.M. and F.S.D. contributed new analytic pipelines; S.S. and J. Heitman supervised the study; J. Huang., C.J.L., S.J.P., V.Y., P.M.M., S.S., and J. Heitman wrote the paper.

