## Supporting Information for "Distinct evolutionary trajectories following loss of RNA interference in *Cryptococcus neoformans*"

**This PDF file includes:**

Supporting text
Figures S1 to S13
Tables S1 to S7
Legends for Datasets S1 to S15
SI References

### Supporting text

### Extended Materials and Methods

#### Strains and growth

The *C. neoformans* and *C. deneoformans* strains analyzed in this study are listed in SI Appendix, Table S6. Strains were stored at -80°C and supplemented with 15% glycerol for long-term storage. Strains were inoculated on YPD solid medium at 30°C for 3 days and maintained at 4°C no more than two weeks. Fresh cells were streaked out from glycerol stocks as needed.

#### Pipeline constructions

Construction of reference-based genome assemblies for Pipeline 1 (Neostop). Raw Illumina sequencing reads generated by Desjardins et al. 2017 were downloaded from the NIH Sequence Read Archive (BioProject ID PRJNA382844) (1). For each of the 387 BioSamples (corresponding strains of interest) associated with the BioProject, a reference-based genome assembly was generated by aligning paired-end sequence data to the genome of the *C. neoformans* reference strain H99 (FungiDB R53) in Pipeline 1. In cases where there were multiple sequencing runs for a given BioSample, the sequencing run containing the largest number of paired-end reads was used. To produce reference-based genome assemblies, reads were aligned to the reference genome using BWA (v0.7.12), variants called using FreeBayes (v1.1), and strain-specific consensus assemblies generated by instantiating the called variants onto the reference genome (2, 3) The read alignment, variant calling, and consensus assembly were carried out using the Snippy (https://github.com/tseemann/snippy) pipeline tool. Variant sites were called with a minimum read depth of 10, a minimum read mapping quality of 60, and a stringent variant quality score of 100 (Phred-scale). This choice of parameters settings produces very conservative variant calls. Following construction of consensus assemblies, genome feature annotation was "lifted over" from the H99 reference genome to each strain-specific genome using the software tool Liftoff (v1.6.3) (4). The `-polish` option of Liftoff was employed to re-align exons in cases where the lift-over procedure resulted in start/stop codon loss or introduced an in-frame stop codon. Based on the polished lift-over annotation, the AGAT GTF/GFF Toolkit software (https://github.com/NBISweden/AGAT) was used to predict protein sequences for all annotated genes in each strain-specific assembly using the `agat_sp_extract_sequences.pl` script. Where multiple protein isoforms are annotated in the reference genome, predictions were generated for each isoform. For each RNAi-related gene of interest (SI Appendix, Table S1), the predicted amino acid sequences of each of the 387 *C. neoformans* strains was compared to the H99 reference genome (FungiDB R53) with Pipeline 1. Candidate loss-of-function alleles were classified as those encoding proteins whose predicted length is < 90% of the model protein length estimated from the entire set of strains. All seven predicted variants by Neostop were validated through Sanger sequencing.

A second, independent program to identify mutations of interest in *C. neoformans* sequencing data was constructed. Polymorphisms were independently identified with the GenesComp program (Pipeline 2), which is available through GitHub (https://github.com/derf0/genescomp). This is a species-specific program for finding polymorphisms by combining multiple factors for evaluating genomic variation. GenesComp attempts to classify polymorphisms according to their likelihood to alter function of the gene product. This includes polymorphisms that may increase the amount of protein, such as changes from rarely used codons to more frequently used codons, as well as changes in introns from poor splice site or branch site sequences to more advantages sequences. For *C. neoformans*, this analysis was carried out with the *C. neoformans* strain H99 (5) as the reference, and intron branch site, splice site, and permissible length rules determined for this strain, as well as codon usage (GenBank: CP003820 through CP003834). Gain of stop codons (nonsense mutations) and loss of stop codons ("resense" mutations) were detected, along with insertions/deletions, in-frame and out-of-frame, and synonymous and non-synonymous polymorphisms. Non-synonymous polymorphisms were categorized as either conservative or non-conservative. Polymorphisms were also identified in UTR regions and in RNA-coding genes such as those encoding tRNAs. The program uses the genomic sequence and annotation of the reference genome, so the accuracy of the predictions is dependent on the accuracy of the annotations and reference strain genome assembly. Sequencing errors were addressed by first performing *de novo* assembly to identify genomic features, followed by aligning reads using BWA/SAMtools to assess base quality scores from the aligned reads through manual curation. Notably, eight out of nine identified variants have been verified by Sanger sequencing, demonstrating the robustness of the approach. The program is written in Python3, though uses several external programs: EMBOSS package version 6 of DNA/RNA/protein programs, Blastx NCBI stand-alone blast package, search Smith-Waterman alignment, and tRNAscan-SE (6–9).

**Small RNA isolation, sequencing, and analysis**

Briefly, strains were cultured in 8 mL YPD liquid medium overnight at 30°C roller drum with 70 rpm. Overnight cultures were pelleted and frozen at -80°C for 1 h. Cells were then lyophilized overnight. Small RNA was isolated from ~70 mg of the lyophilized cell using the mirVana™ miRNA Isolation Kit (Thermo Fisher) following manufacturer's instructions. Small RNA sequencing libraries were prepared using QIASeq miRNA Seq Auto Kit (Qiagen) and sequenced on the Illumina NextSeq 500 (1x75 bp) or Novaseq X Plus (2x150 bp) platforms at the Duke University Sequencing and Genomic Technologies Core facility. Sequencing adaptors were removal by Cutadapt (v1.18) (10), followed by discarding untrimmed reads or reads shorted than 14 nt or longer than 40 nt. The remaining reads were mapped to appropriate reference genome with Bowtie (v1.3.1) (11) that allowing multiple alignments (Bt65 and XL280α) or up to five alignments (H99), but no mismatches. Custom perl script was used for counting read size distribution and 5'-nucleotide identities from 14 to 28 nt (https://github.com/timdahlmann/smallRNA) (12). The sorted and indexed BAM file, generated after bowtie mapping was used as input for counting and visualizing the normalized read coverage (RPM, Read Per Million) in the coordinates of specific transposon element using sRNA_Viewer (https://github.com/MikeAxtell/sRNA_Viewer). All sRNA analyses were performed in at least two biological replicates, except for Bt52, Bt152, Bt208, Bt210, and LP-RSA2296 (one biological replicate).

**Fluctuation assay and mutational spectra analysis**

Briefly, 10 to 20 independent cultures of each strain were grown in 5 mL YPD for 12 to 18 h in a roller drum set to 70 rpm and at 30°C. The cultures were then centrifuged at 3000xg for 5 min before being resuspended in sterile water. The resuspended cultures were diluted 100,000x, and 100 µL of each were plated on YPD to assess the total number of cells in each culture. 100 µL of the undiluted cultures were plated on YPD medium containing 100 ng/mL rapamycin + 1 µg/mL FK506 (R+F) or YNB containing 1 mg/mL 5-Fluoroorotic Acid (5-FOA). For assessing hypermutation on YNB medium containing 100 µg/mL

5-fluorocytosine (5-FC), the resuspended cultures were diluted 100x, and 100 µL were plated. Colonies were counted following incubation for 3 to 4 days on YPD at 30°C and 6 to 12 days on drug plates (30°C for 5-FOA and 5-FC, and 37°C for R+F) using an eCount colony counter and ImageJ.

To ensure the identified mutant events were independent, only one resistant colony per drug plate was purified, as described previously (20). DNA extracted from independent drug-resistant colonies was used for genotyping PCR of target genes (e.g., *FRR1*, *FUR1*, *UXS1*, and *FCY2*). *ACTIN* was amplified as a DNA quality and loading control. LaTaq was utilized for amplifying extra-long PCR products (NEB). The PCR products were sequenced using Sanger or Plasmidsaurus amplicon sequencing (https://www.plasmidsaurus.com/), allowing for the detection of sequence variations.

**Nanopore long-read sequencing and *de novo* genome assembly**

High molecular-weight DNA was extracted by CTAB method as described previously (13). DNA size was estimated by pulsed-field gel electrophoresis and quantified by Qubit high sensitivity assay kit. Nanopore Sequencing library was prepared by using native barcoding kit SQK-NBD114.24 following manufacturer's protocol. Up to 2 samples were barcoded and sequencing in R10.4.1 flow cells (FLO-MIN114) for 72h. After sequencing, raw Pod5 files were trimmed, basecalled and demultiplexed with Dorado (v0.5.3, https://github.com/nanoporetech/dorado). Canu (v2.2) (14) was used for *de novo* genome assembly following sequence correction with medaka (v1.11.3, https://github.com/nanoporetech/medaka) and up to 5 rounds of Pilon (v1.24) (15) polish with Illumina WGS reads. Naming and orientation of each contigs were adjusted based on the H99 reference genome (https://www.ncbi.nlm.nih.gov/datasets/genome/GCF_000149245.1/). Unplaced contigs with >50% duplicated with 14 core chromosomes or bacterial DNA were removed from the assemblies. Telomere repeats (5' (TAACCCC)n '3) were found on both ends of the 14 chromosomes of Bt210 and LP-RSA2296 in the assemblies. The repeat content and annotation analysis were performed using RepeatMasker (v4.0.7) with Dfam (v3.3) and RepBaseRepeatMaskerEdition-20181026 libraries that was supplemented with RepBase EMBL database (v26.04). All the repeats identified were used for the comparison between three strains, H99, Bt210 and LP-RSA2296.

### CRISPR-mediated allele exchange and gene deletion

Guide RNAs (gRNAs) for allele exchange assays were designed by using EuPaGDT website (http://grna.ctegd.uga.edu/) and constructed as previously described (16, 17). Donor DNA containing the sequence of interests was generated by mutating the endogenous sequences via Q5 Site-Directed Mutagenesis Kit (for causing the *rde4* splice-site mutation or the *znf3* non-sense mutation in Bt65+*ZNF3*) (NEB). Safe Haven 1 (SH1) locus was used for integration in transformants (18).

Strains were transformed with 500 ng *C. neoformans* codon optimized Cas9 (17), 2 μg allele exchange donor DNA and 300 ng gRNA against the mutation of interest as well as 300 ng gRNA targeting SH1 and 2 μg AscI digested pSDMA25 plasmid (18), via a transient CRISPR-Cas9 coupled with electroporation (TRACE) system as previously described (19). Potential transformants were selected on YPD+NAT, DNA extracted from resistant colonies were used as template for amplifying a region outside the allele swap donor for avoiding the condition of ectopic insertion. In addition to the desired mutations, no additional mutations were found by amplicon sequencing (Plasmidsaurus). The transformants with expected sequence alternation were further validated for the presence of Cas9, gRNA cassette and the status for SH1 by genotyping PCR.

To delete the *RDP1* gene in *C. deneoformans* 37-02, 1,069 bp upstream and 1,031 bp downstream flanking sequences of *RDP1* were fused to *NAT* marker amplifying from pAI3 plasmid by split-marker method. Two independent gRNAs were designed for improving knockout efficiency. 37-02 was co-transformed with Cas9, dual gRNAs, and the *NAT* split marker by the TRACE protocol (19). Internal, 5' junction and 3' junction PCR were utilized for confirming the successful gene deletion.

### *In vitro* passage assay and preliminary hypermutator screening

Five single colonies of Bt210, LP-RSA2296, and H99 were passaged as patches on YPD medium at 30°C and 37°C for 28 and 56 passages, respectively, every 48 hours. Two independent cultures per strain were cultured overnight at 30°C and resuspended in 2 mL sterilizer water. Homogenized cultures after vortex were swab on a quadrant of the YPD medium containing 100 ng/mL rapamycin + 1 μg/mL FK506 (R+F).

**Phylogenetic analysis**

The sequences of identified KDZ-transposase-domain containing proteins from *Coprinopsis cinerea* were obtained from NCBI GenBank (https://www.ncbi.nlm.nih.gov/genbank/) (21). The maximum-likelihood phylogeny was constructed with MEGA X (v10.0.1) with 1000 bootstrap replicates (22).

**Mating and spore dissection**

A cross between Bt210 and Bt65 was performed as previously described (23). A total of 57 F1 progeny were isolated via random spore microdissection from the MS plate, which were plated with a mixed cell suspension containing both Bt210 and Bt65 (23).

**Transposon copy number estimates from assembled genomes using BLAT**

To estimate transposon copy number from assembled genomes, we used BLAT (24) to search for regions homologous to KDZ1, KDZ2, KDZ3, and Cnl1 in the assembled *C. neoformans* genomes given in SI Appendix, Dataset S15. The sequences of each transposon were taken from the Bt65 genome where the KDZ transposons were first identified. BLAT searches were run with a minimum identity of 0.9. The BLAT accessory tools pslReps was subsequently used to find the best alignments for each query sequence with a minimum coverage of 0.95. The number of such best alignments per genome of interest was used as the BLAT-based estimate of transposon copy number.

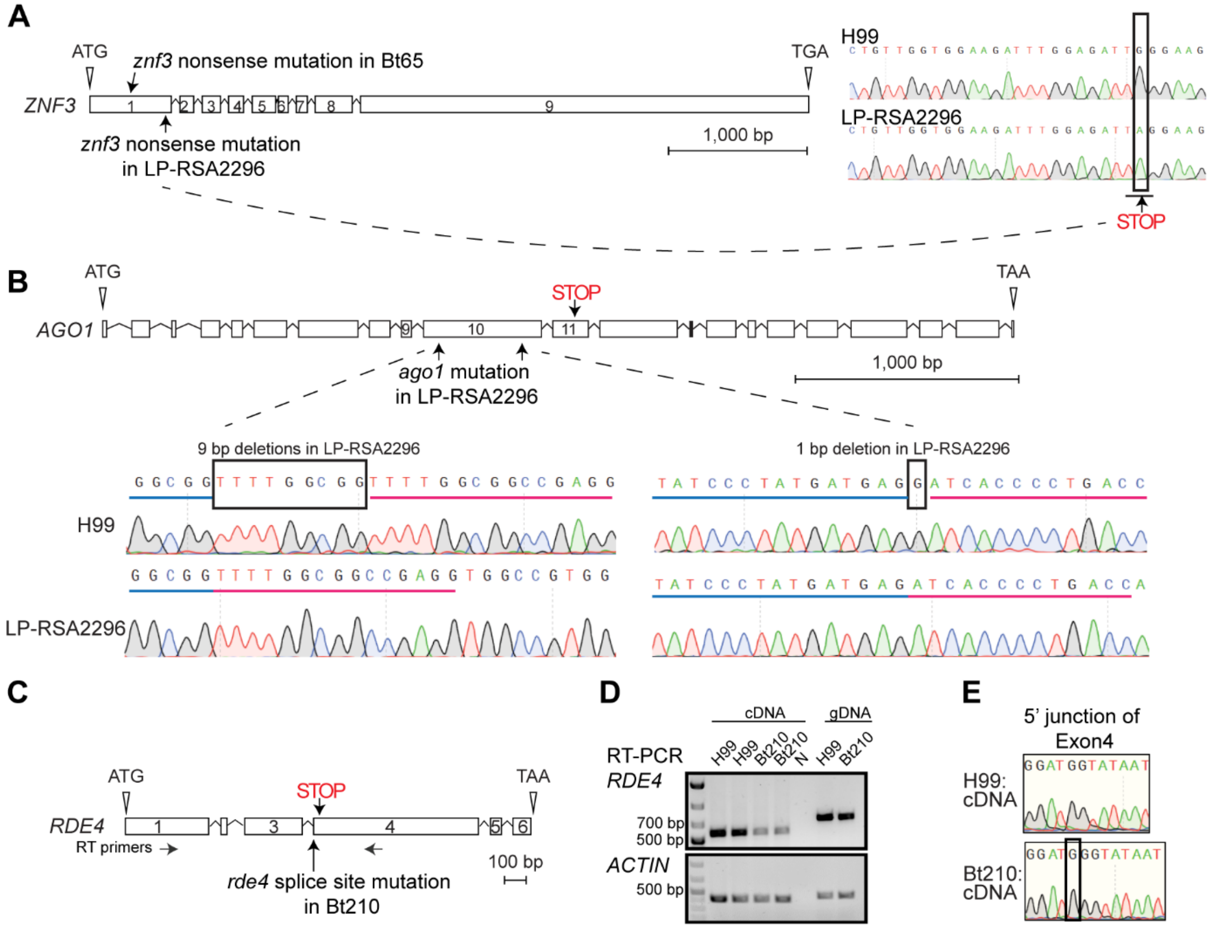


**Fig. S1. Confirmation of the identified mutations in LP-RSA2296 and Bt210.**
(A) Schematic diagram of the *ZNF3* gene structure. The numbers within the *ZNF3* gene structure indicate the order of the *ZNF3* exons. The positions of *ZNF3* mutations found in Bt65 and LP-RSA2296 are labeled. The *ZNF3* nonsense mutation predicted in LP-RSA2296 was confirmed with Sanger sequencing and is indicated by a black box. (B) Schematic diagram of the *AGO1* gene structure. The numbers within the *AGO1* gene structure indicate the order of the *AGO1* exons. The predicted deletions in *AGO1*, leading to the protein truncation in exon11, were confirmed with Sanger sequencing. The identical sequences surrounding the deletions are highlighted with blue and red lines. (C) Schematic diagram of the *RDE4* gene structure. The numbers within the *RDE4* gene structure indicate the order of the *RDE4* exons. Arrows below the gene structure represent the binding sites of primers used for *RDE4* RT-PCR. (D) RT-PCR analysis of the *RDE4* splicing error. Two independent cDNA amplifications were performed for each sample. Genomic DNA was used to rule out the possibility of gDNA contamination in the cDNA. *ACTIN* amplification served as a loading control. Lane 'N' indicates the water negative control. (E) Confirmation of the *rde4* splicing error by Sanger sequencing. *RDE4* RT-PCR products from H99 and Bt210 cDNA were subjected to Sanger sequencing. An additional G caused by the splicing error near the 5' junction of exon4 is highlighted by a black box.

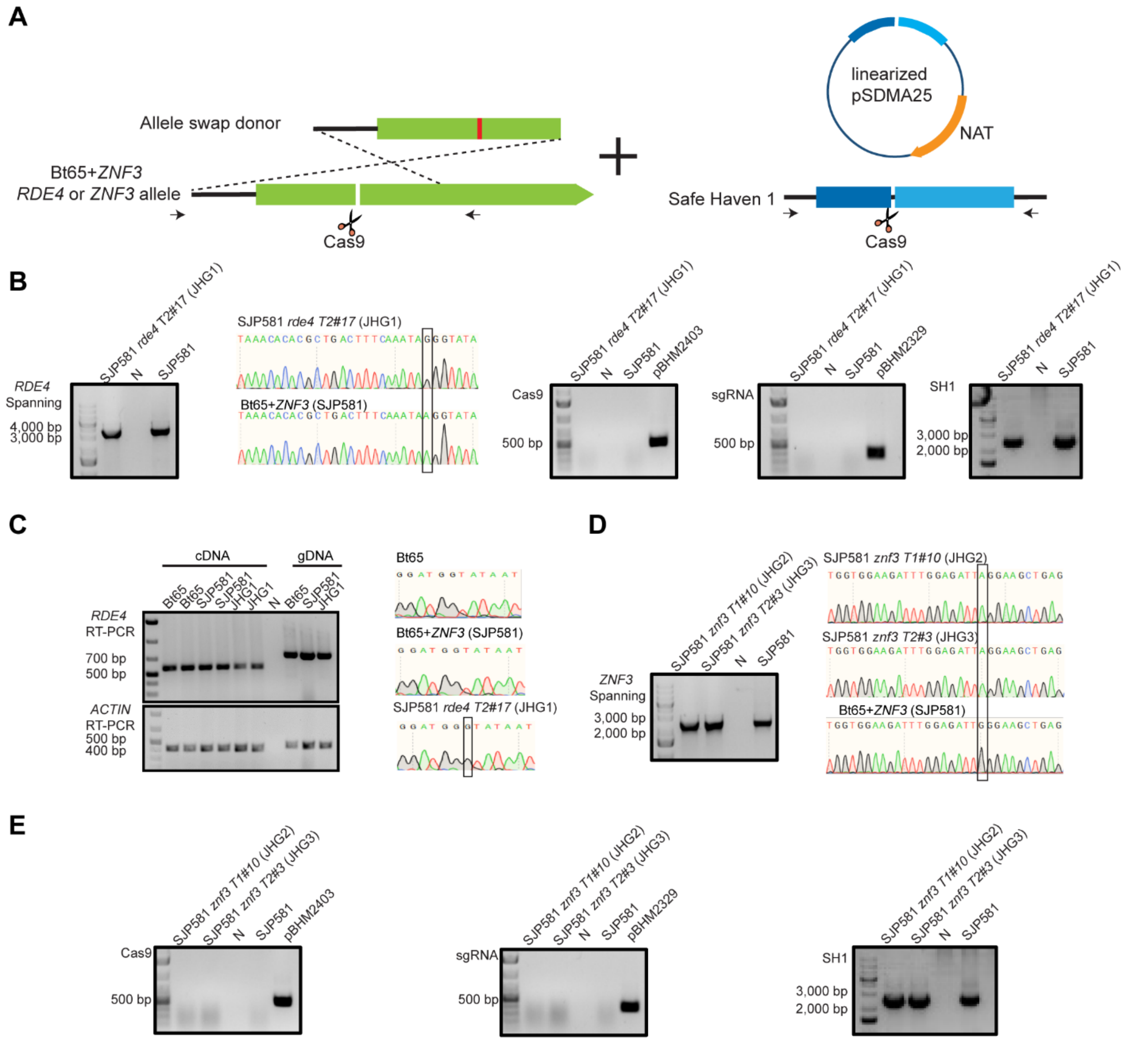


**Fig. S2. Confirmation of transformants from *rde4* and *znf3* CRISPR-mediated allele exchange assay.** (A) Schematic diagram of the CRISPR-mediated allele exchange assay. An allele exchange donor containing only the desired mutation (red bar) was constructed to introduce the mutation in either *RDE4* or *ZNF3* in the Bt65+*ZNF3* (SJP581) background to repair the DNA double-strand break near the sequence of interest (white bar) caused by Cas9. After Cas9 digestion, the Safe Haven 1 locus was used for integrating the linearized pSDMA25, which contains a nourseothricin (NAT) resistance selectable marker. Arrows under the *RDE4* or *ZNF3* alleles and Safe Haven 1 indicate the position of genotyping primers. Light and dark blue regions in the plasmid pSDMA25 indicate the homologous sequences used for recombination. The NAT resistance selectable marker is labeled in orange. (B) Primers spanning the *RDE4* locus, as indicated in (A), were used for genotyping Bt65+*ZNF3 rde4*. Lane 'N' indicates a water negative control, while genomic DNA from SJP581 served as the positive control. The purified products were sequenced by amplicon sequencing (Plasmidsaurus) and Sanger sequencing. The desired mutation is highlighted (black box) in the Sanger sequencing results. Genotyping PCRs were performed for Cas9, the gRNA cassettes, and the *SH1* locus. No stable Cas9 or gRNA cassettes were found in the transformants, and no NAT marker integration was found in the SH1. Plasmid DNA from plasmids pBHM2403 and pBHM2329 served as positive controls for amplifying Cas9 and the gRNA cassettes, respectively. (C) RT-PCR for the *RDE4* locus was performed for Bt65, Bt65+*ZNF3*, and Bt65+*ZNF3 rde4* cDNA and gDNA. Lane 'N' indicates the water negative control. Two independent cDNA amplifications were performed for each sample. Amplified products from cDNA were purified and used for Sanger sequencing. An additional 'G' frameshift caused by the splicing error is highlighted in a black box. (D and E) A similar genotyping strategy as in (B) was applied to obtain two independent Bt65+*ZNF3 znf3* transformants.

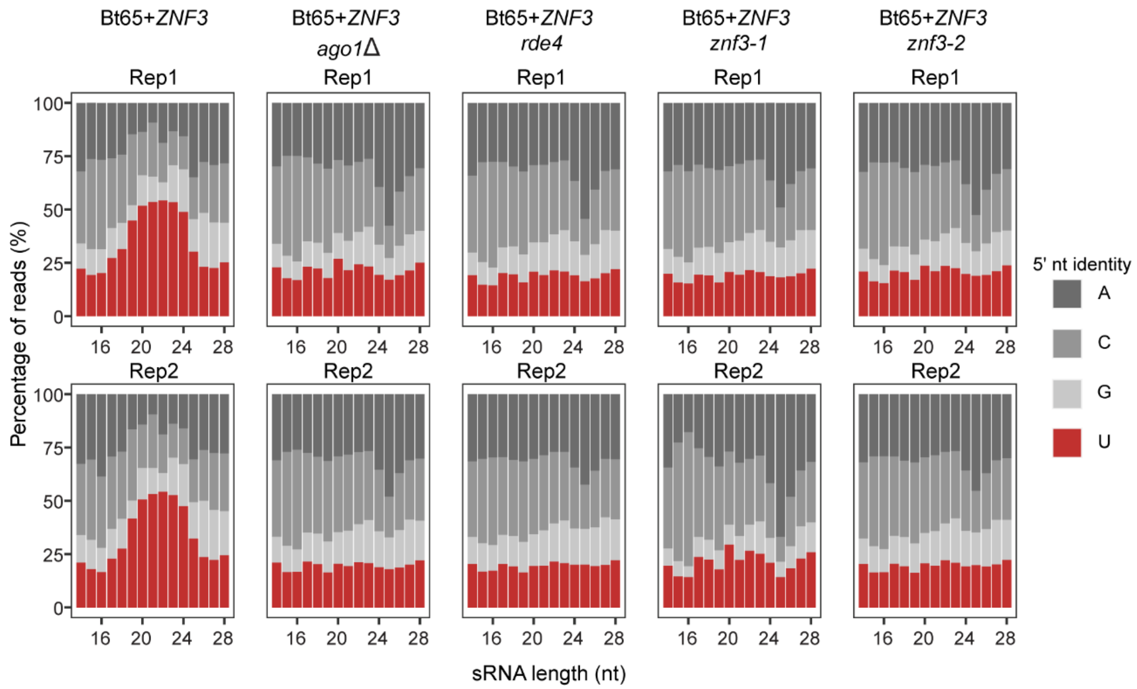


**Fig. S3. Proportion of 5'-nucleotide identity in the sRNA reads from the allele exchange strains.** The upper and lower panels are from two independent biological replicates (Rep1 and Rep2). Bt65+*ZNF3* and Bt65+*ZNF3 ago1*Δ served as positive and negative controls, respectively. All of the sRNA reads were mapped to the Bt65 genome.

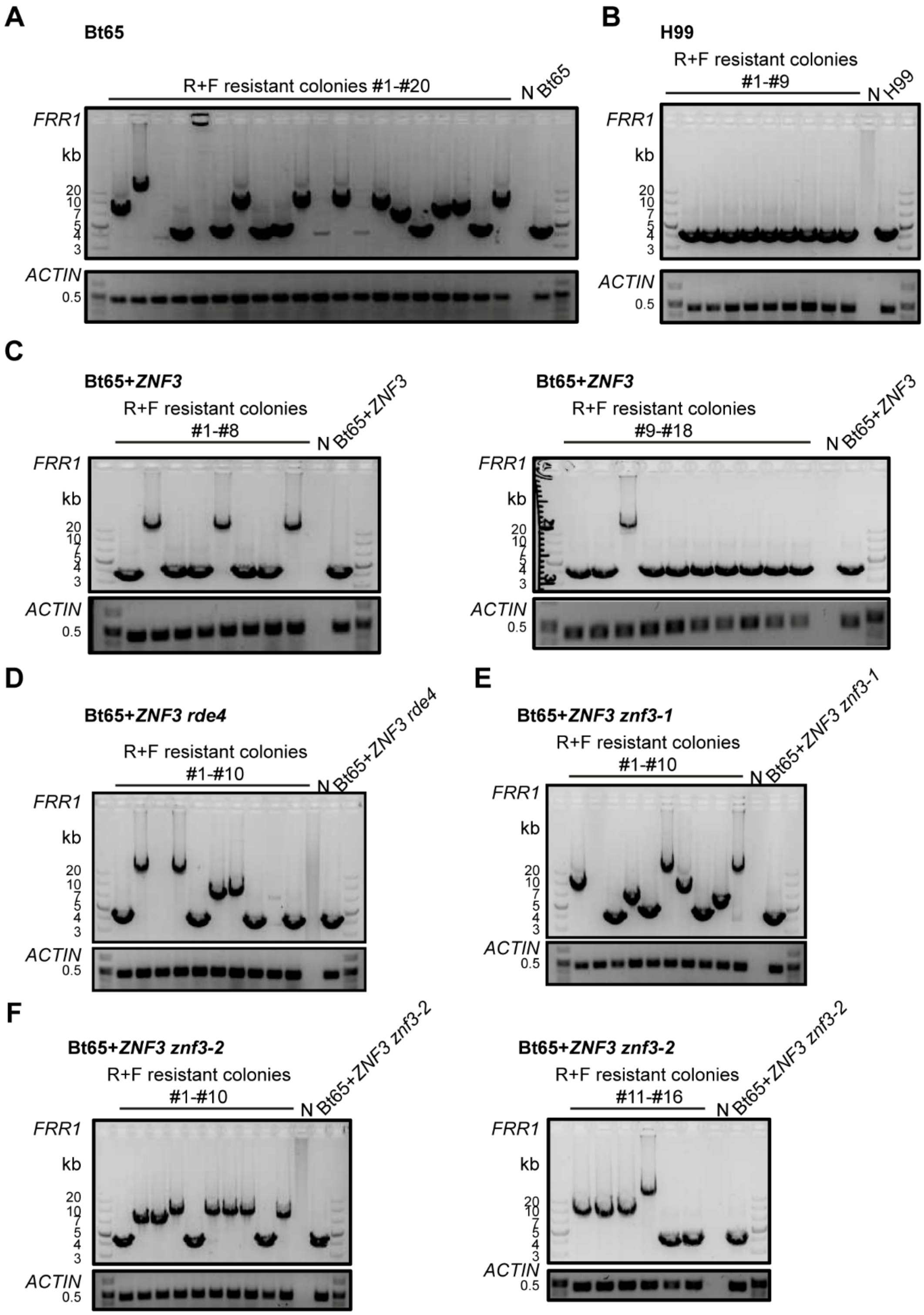


**Fig. S4. Gel electrophoresis of *FRR1* products from independent R+F resistant colonies generated in allele exchange assay.**

Representative gel images of *FRR1* PCR products of R+F resistant colonies from (A) Bt65, (B) H99, (C) Bt65+*ZNF3*, (D) Bt65+*ZNF3 rde4*, (E) Bt65+*ZNF3 znf3-1*, and (F) Bt65+*ZNF3 znf3-2*. Lane 'N' indicates the water negative control. PCR amplification of *FRR1* in the R+F sensitive progenitor produces a ~3.7kb electrophoretic species (primers JH60/61).

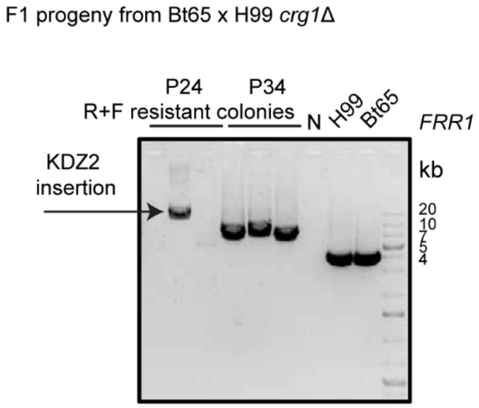


**Fig. S5. KDZ2-mediated R+F resistance in Bt65 derived F1 progeny.**
Gel image of *FRR1* PCR products of R+F resistant colonies from Bt65 x H99 *crg1*Δ F1 progeny P24 and P34. The colony with a KDZ2 insertion is labeled. Lane 'N' indicates the water negative control. PCR amplification of *FRR1* in the R+F sensitive parents produces a ~3.7 kb electrophoretic species (primers JH60/61).

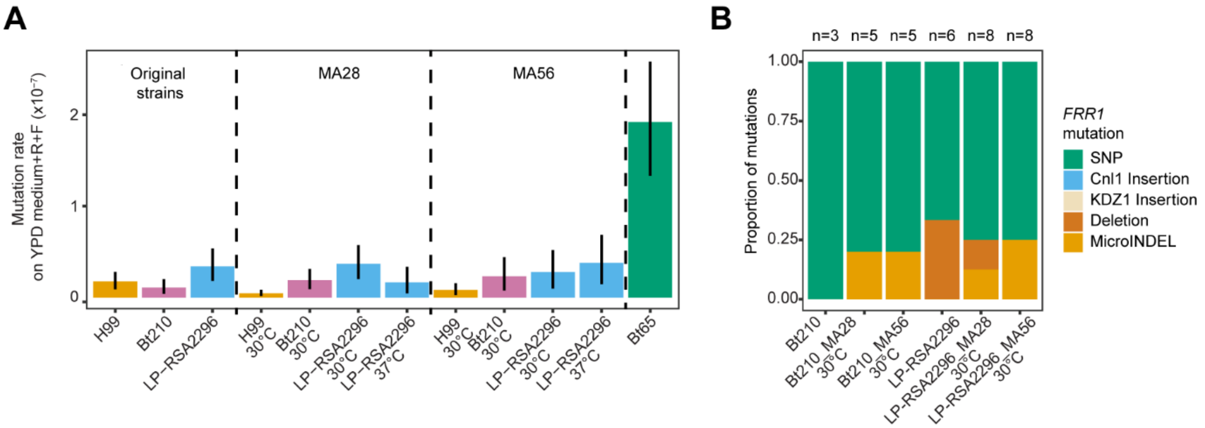


**Fig. S6. Short-term experimental evolution in Bt210 and LP-RSA2296.**
(A) Mutation rates of the progenitor strains H99, Bt210, LP-RSA2296, and their derived strains after 28 (MA28) or 56 (MA56) *in vitro* passages, when grown on YPD medium containing rapamycin + FK506, with the hypermutator Bt65 serving as positive control. The error bars represent 95% confidence intervals. (B) Mutational spectra in the *FRR1* gene of the independent R+F resistant colonies from each indicated strain.

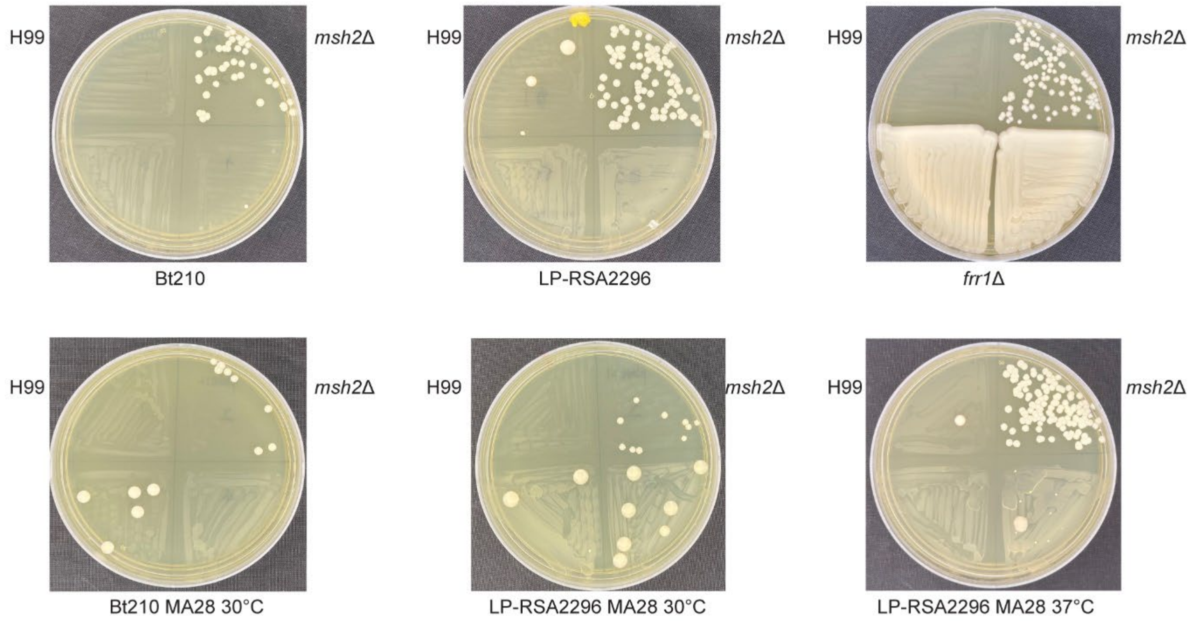


**Fig. S7. Preliminary hypermutator screening**.
Two independent overnight cultures of Bt210, LP-RSA2296, Bt210 MA28 30 °C, LP-RSA2296 MA28 30 °C and LP-RSA2296 MA28 37 °C were swabbed on a single quadrant of a plate containing YPD medium+R+F for hypermutator screening. H99 served as the negative control, while *msh2*Δ and *frr1*Δ served as the positive controls for screening. Colonies were photographed 7 days post-plating.

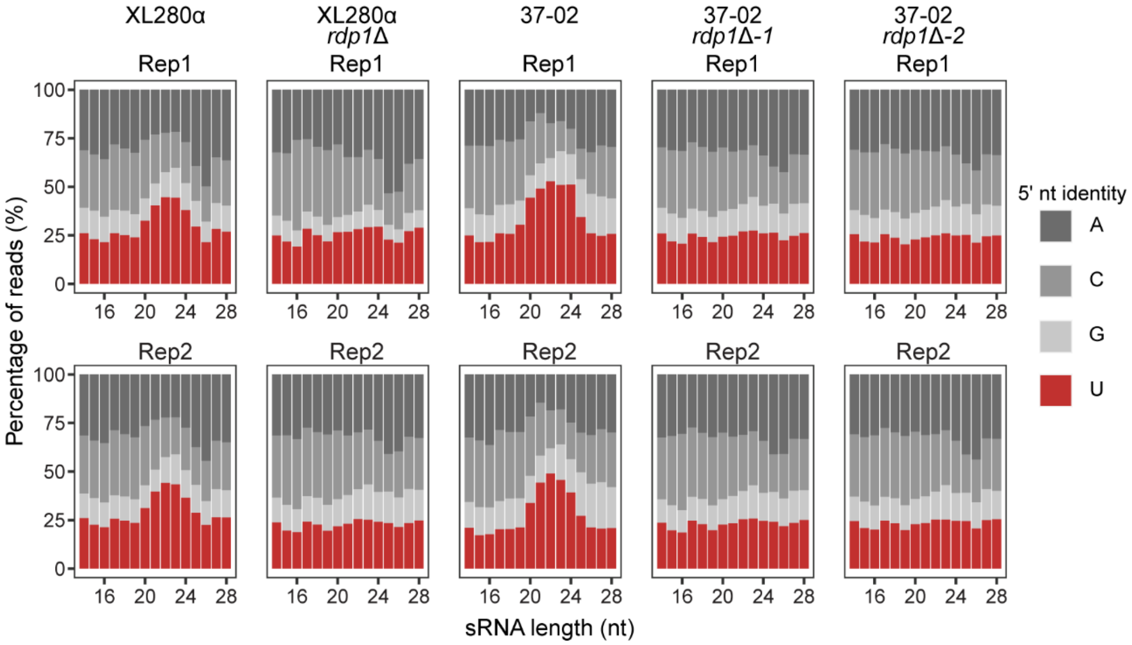


**Fig. S8. Proportion of 5'-nucleotide identity in sRNA reads from the XL280α derived strains.** The upper and lower panels are from two independent biological replicates (Rep1 and Rep2). XL280α and XL280α *rdp1Δ* served as positive and negative controls, respectively. All of the sRNA reads were mapped to the XL280α genome.

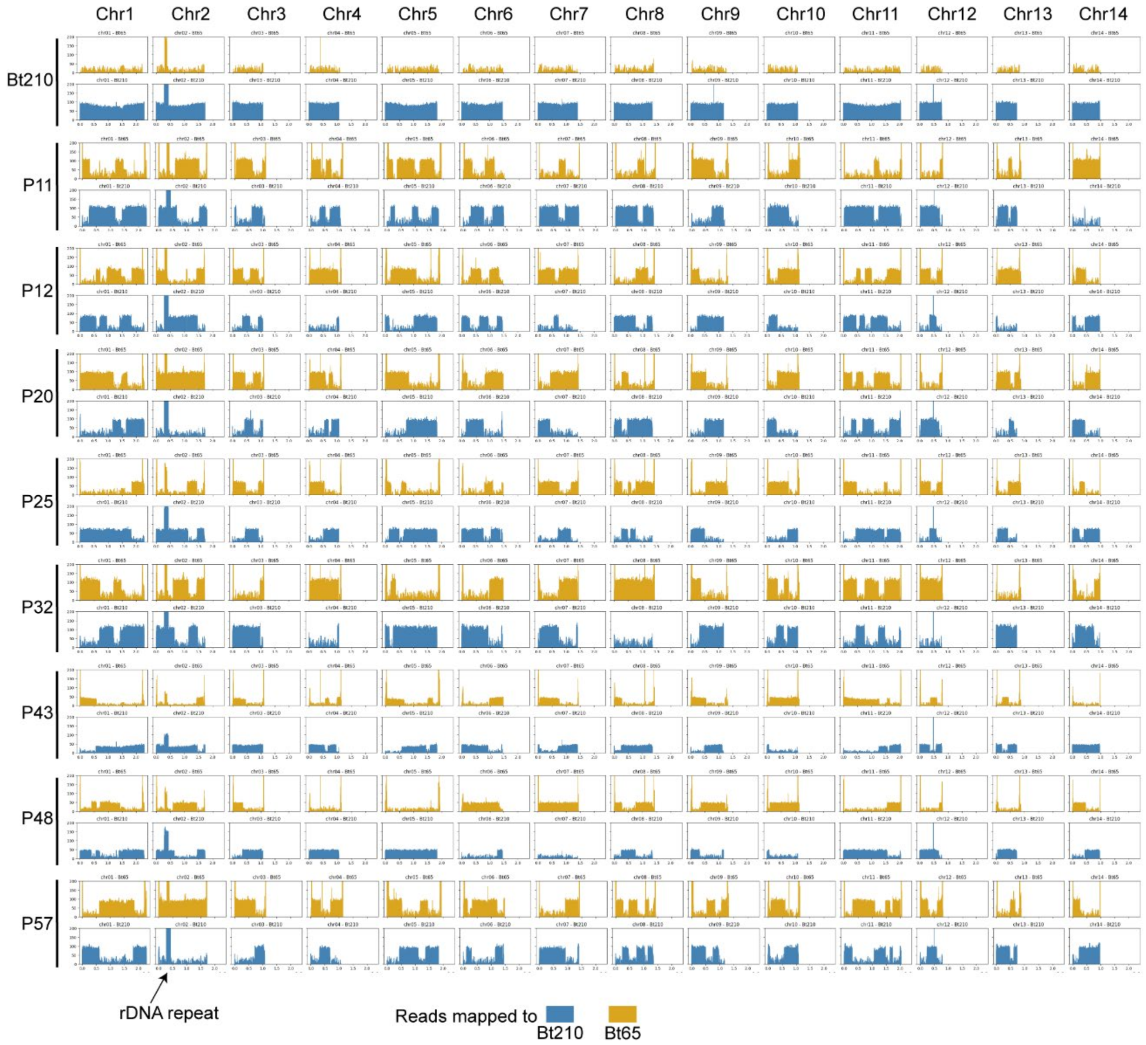


**Fig. S9. Recombination map of the F1 progeny from the Bt65a x Bt210α cross.**
Whole-genome sequencing reads from 8 F1 progeny were mapped to a concatenated Bt65**a**/ Bt210α reference genome. Recombination was inferred based on the relative depth against the two references (Orange-Bt65 and Blue-Bt210). The mapping result from Bt210 served as a control for the background level of reads mapping. Please note that the x-axis of all plots has been set to the length of chromosome 1, and the boundaries of all of the other chromosomes can be inferred by the ends of their respective mapping plots.

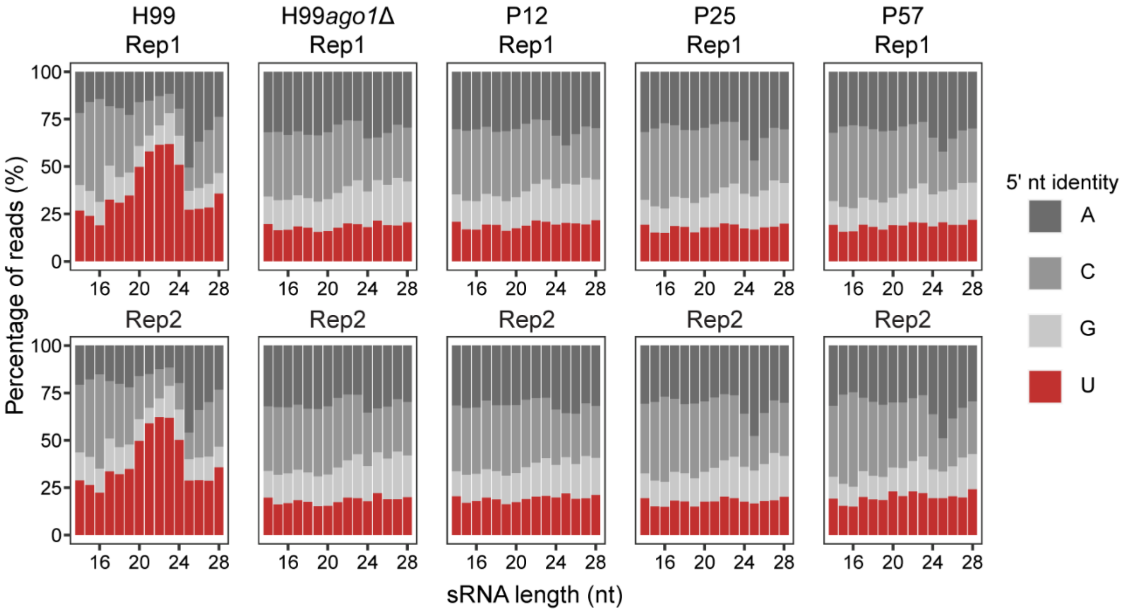


**Fig. S10. Proportion of 5'-nucleotide identities in sRNA reads from the F1 hypermutator progeny.** The upper panel and the lower panels are from two independent biological replicates (Rep1 and Rep2). H99 and H99 *ago1*Δ served as positive and negative controls, respectively. All the sRNA reads were mapped to the Bt65 genome.

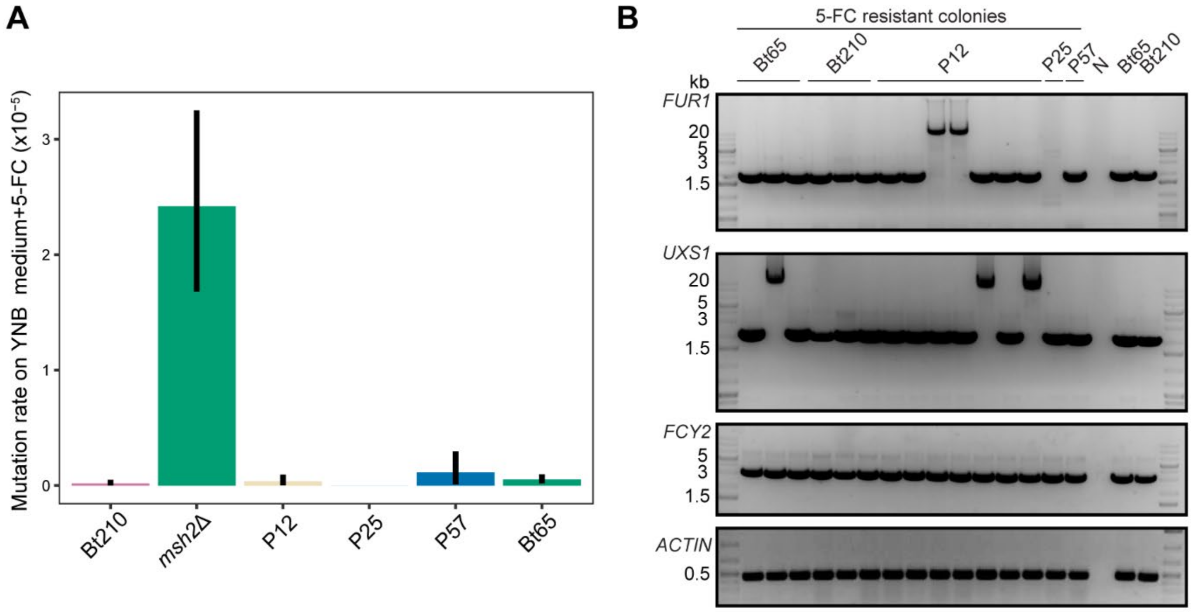


**Fig. S11. Insertion of KDZ1 leads to 5-FC resistance.**
(A) Mutation rates of three F1 hypermutator progeny when grown on YNB medium containing 5-fluorocytosine (5-FC), with parental strains Bt210, Bt65, and *msh2*Δ serving as negative and positive controls, respectively. The error bars represent 95% confidence intervals. (B) Genotyping PCR for independent 5-FC resistant colonies. Three known targets of 5-FC (*FUR1*, *UXS1*, and *FCY2*) were selected for genotyping, with *ACTIN* serving as a DNA quality and loading control. Lane 'N' indicates the water negative control.

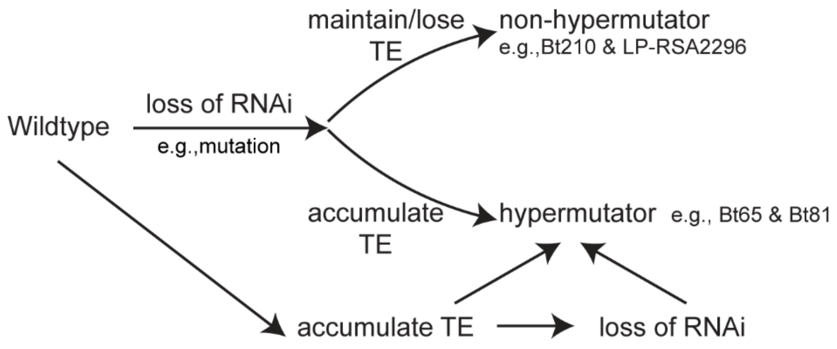


**Fig. S12. Current model for paths to hypermutation.**
After RNAi loss, isolates (e.g., Bt65 and Bt81) underwent transposon accumulation and evolved into hypermutators. In contrast, some other RNAi-deficient isolates (e.g., Bt210 and LP-RSA2296) might maintain or lose transposons and remain non-hypermutators. Additionally, transposon accumulation might occur without RNAi loss and cause a hypermutator phenotype with or without subsequent RNAi loss.

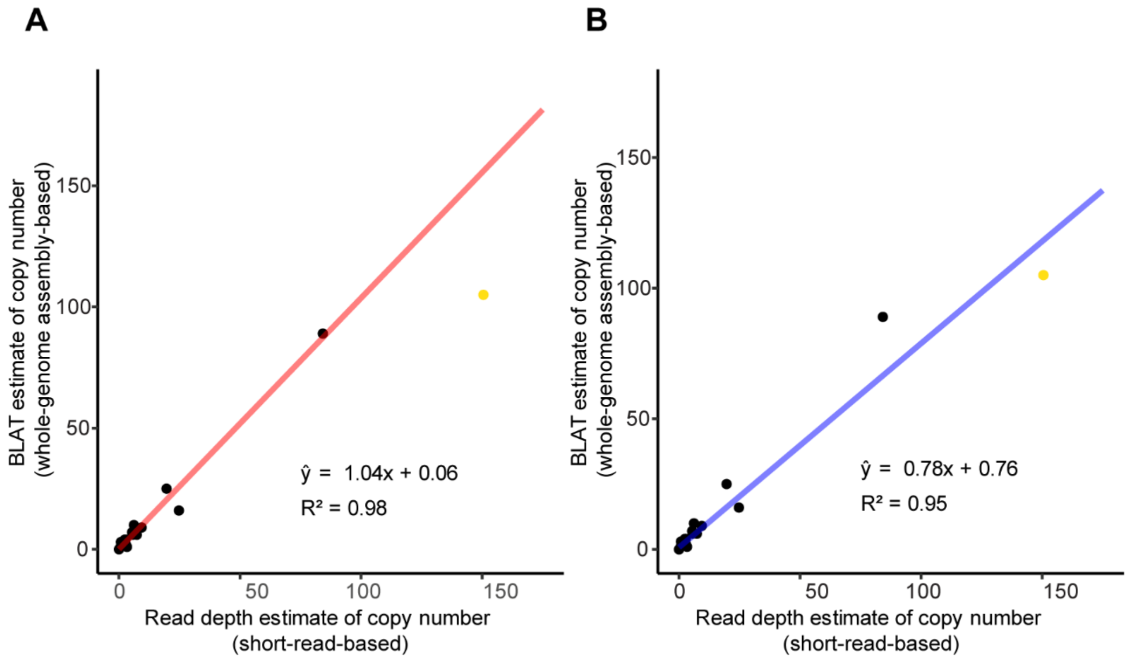


**Fig. S13. Correlation between the estimated transposon copy numbers from whole-genome assembly-based and short-read-based methods.**
Linear regression was used to estimate the relationship between different transposon (KDZ1, KDZ2, KDZ3, and CnI1) copy number estimations (whole-genome assembly-based method vs. short-read-based method). The CnI1 copy number in Bt65 (yellow dot) was excluded in (A) but included in (B). The whole-genome assemblies used are listed in Dataset S15.

**Table S1. Genes required for RNAi in *Cryptococcus*.**

| Gene name[1] | Gene ID | Encoded function[2] | References |
|---|---|---|---|
| *AGO1* | CNAG_04609 | Argonaute; required for silencing during MIS and SIS | (25, 26) |
| *RDP1* | CNAG_03466 | RNA-dependent RNA polymerase; required for silencing during MIS and SIS | (25) |
| *QIP1* | CNAG_01423 | An ortholog of *Neurospora crass* exonuclease: Quelling-defective-2 interacting protein; required for MIS & SIS | (27, 28) |
| *ZNF3* | CNAG_02700 | Zinc finger protein; Required for MIS & SIS | (28, 29) |
| *GWC1* | CNAG_06486 | GW/WG protein; Required for MIS; role in SIS unknown | (27) |
| *DBR1* | CNAG_03222 | Lariat debranching enzyme; required for MIS; role in SIS unknown | (27) |
| *RPA70* | CNAG_01144 | Replication protein A subunit; required for MIS; role in SIS unknown; essential for viability | (30) |
| *RPA32* | CNAG_01316 | Replication protein A subunit; required for MIS; role in SIS unknown; mating defects in deletion mutants | (30) |
| *RDE1* | CNAG_01848 | *R*NAi-*De*fective protein 1; G-patch protein involved in rRNA maturation & splicing; Required for MIS; role in SIS unknown | (31) |
| *RDE3* | CNAG_06643 | *R*NAi-*De*fective protein 3; non-Dicer RNase III; required for MIS; role in SIS unknown | (31) |
| *RDE4* | CNAG_01157 | *R*NAi-*De*fective protein 4; putative terminal nucleotidyltransferase; required for MIS; role in SIS unknown | (31) |
| *RDE5* | CNAG_04791 | *R*NAi-*De*fective protein 5; Required for MIS; role in SIS unknown | (31) |

[1]: Genes that are involved but not required for RNAi in *C. neoformans* (e.g., *DCR1* (CNAG_02742)(25), *DCR2* (CNAG_02745) (25), and *CPR2* (CNAG_03938)(28)) have not be included in this list
[2]: MIS: mitotic-induced silencing; SIS: sex-induced silencing

**Table S2. Identified KDZ transposase domain-containing proteins in the H99 and Bt65 genomes.**

| Isolate | Description[1] | Full sequence E-value[2] | Note[3] |
|---|---|---|---|
| H99 | CNAG_00128 | 1.1e-48 | KDZ1 |
| H99 | CNAG_06757 | 3.3e-48 | |
| H99 | CNAG_04102 | 2e-22 | |
| H99 | CNAG_07523 | 2.8e-07 | |
| H99 | CNAG_02058 | 0.00015 | |
| Bt65 | Bt65_Chr07_000348.1 | 8.6e-52 | KDZ2 |
| Bt65 | Bt65_Chr05_000319.1 | 7.80E-48 | KDZ1 |
| Bt65 | Bt65_Chr02_000042.1 | 6.60E-47 | KDZ-containing protein 5 |
| Bt65 | Bt65_Chr09_000343.1 | 7.30E-42 | KDZ1 |
| Bt65 | Bt65_Chr06_000122.1 | 8.30E-42 | KDZ1 |
| Bt65 | Bt65_Chr04_000295.1 | 8.40E-42 | KDZ1 |
| Bt65 | Bt65_Chr07_000538.1 | 8.40E-42 | KDZ1 |
| Bt65 | Bt65_Chr04_000319.1 | 3.90E-38 | KDZ1 |
| Bt65 | Bt65_Chr09_000136.1 | 6.80E-32 | KDZ3 |
| Bt65 | Bt65_Chr01_000480.1 | 4.10E-21 | KDZ-containing protein 4 |
| Bt65 | Bt65_Chr09_000017.1 | 8.00E-21 | KDZ-containing protein 6 |

[1]: The H99 annotation was obtained from (https://www.ncbi.nlm.nih.gov/datasets/genome/GCF_000149245.1/) and the Bt65 annotation was predicted by Helixer (https://github.com/weberlab-hhu/Helixer) (5, 32)
[2]: E- values were generated by performing an hmmsearch against the Kyakuja-Dileera-Zisupton transposase domain (PF18758)
[3]: There are 6 copies of KDZ1 in the Bt65 genome

**Table S3. Estimated full-length Cnl1, KDZ1, KDZ2, and KDZ3 copy numbers in the passaged strains.**

| Isolate[1] | Cnl1 | KDZ1 | KDZ2 | KDZ3 |
|---|---|---|---|---|
| H99 | 0 | 1 | 0 | 0 |
| Bt210 | 0 | 1 | 0 | 0 |
| LP-RSA2296 | 0 | 1 | 0 | 1 |
| MA28-H99 30°C | 0 | 1 | 0 | 0 |
| MA28-Bt210 30°C | 0 | 1 | 0 | 0 |
| MA28-LP-RSA2296 30°C | 0 | 1 | 0 | 1 |
| MA28-LP-RSA2296 37°C | 0 | 1 | 0 | 1 |
| MA56-H99 30°C | 0 | 1 | 0 | 0 |
| MA56-Bt210 30°C | 0 | 1 | 0 | 0 |
| MA56-LP-RSA2296 30°C | 0 | 1 | 0 | 1 |
| MA56-LP-RSA2296 37°C | 0 | 1 | 0 | 1 |

[1]: Whole-genome sequences were generated in this study

**Table S4. Copy numbers of full-length transposons of various types in strain XL280α and the passaged strain 37-02.**

| TE | XL280α[1] | 37-02[1] |
|---|---|---|
| Cnl1 | 9 | 49 |
| Tcn12 | 1 | 2 |
| T1 | 5 | 14 |
| T2-short | 3 | 3 |
| T2-long | 4 | 8 |
| T3 | 8 | 7 |
| Crypton Cn1 | 1 | 3 |
| KDZx1 | 1 | 1 |

[1]: Nanopore whole-genome assemblies were obtained from Gusa et al., 2023 (33)

**Table S5. Estimated full-length Cnl1, KDZ1, KDZ2, and KDZ3 copy numbers in the F1 hypermutator and non-hypermutator progeny.**

| Isolate[1] | Cnl1 | KDZ1 | KDZ2 | KDZ3 | Hypermutator when selected on R+F? |
|---|---|---|---|---|---|
| Bt65 | 290 | 4 | 1 | 1 | Yes, Cnl1 hypermutator |
| Bt210 | 0 | 1 | 0 | 0 | No |
| P11 | 185 | 2 | 0 | 1 | No |
| P12 | 188 | 7 | 1 | 0 | Yes, KDZ1 hypermutator |
| P20 | 142 | 2 | 1 | 0 | No |
| P25 | 195 | 3 | 0 | 1 | Yes, Cnl1 hypermutator |
| P32 | 150 | 2 | 1 | 0 | No |
| P43 | 153 | 1 | 0 | 0 | No |
| P48 | 155 | 3 | 1 | 1 | No |
| P57 | 282 | 3 | 1 | 1 | Yes, Cnl1 hypermutator |

[1]: Whole-genome sequences were generated in this study

**Table S6. Strains used in this study.**

| Strain name/designation | Stock number | Genotype | Note | Source |
|---|---|---|---|---|
| H99 | JOHE4413 | *MAT*α | *C. neoformans* natural isolate, VNI lineage | (5) |
| Bt210 | JOHE5170 | *MAT*α | *C. neoformans* natural isolate, VNI lineage | (1, 20) |
| LP-RSA2296 | JOHE22081 | *MAT*α | *C. neoformans* natural isolate, VNI lineage | (1, 20) |
| A2-102-5 | JOHE22038 | *MAT*α | *C. neoformans* natural isolate, VNI lineage | (1, 20) |
| D17-1 | JOHE22039 | *MAT*α | *C. neoformans* natural isolate, VNI lineage | (1, 20) |
| NRHc5028.ENR.STOR | JOHE22792 | *MAT*α | *C. neoformans* natural isolate, VNI lineage | (1, 20) |
| Bt65 | JOHE5067 | *MAT***a** | *C. neoformans* natural isolate, VNBII lineage | (1, 20) |
| Bt81 | JOHE5082 | *MAT***a** | *C. neoformans* natural isolate, VNBII lineage | (1, 20) |
| Bt84 | JOHE5085 | *MAT*α | *C. neoformans* natural isolate, VNBII lineage | (1, 20) |
| Bt52 | JOHE5054 | *MAT*α | *C. neoformans* natural isolate, VNBII lineage | (1, 20) |
| Bt152 | JOHE5148 | *MAT*α | *C. neoformans* natural isolate, VNBII lineage | (1, 20) |
| Bt208 | JOHE5168 | *MAT*α | *C. neoformans* natural isolate, VNBII lineage | (1, 20) |
| H99 *rdp1*Δ | JOHE6840 | *MAT*α *rdp1::NEO* | H99 *rdp1*Δ, RNAi mutant | (34) |
| H99 *ago1*Δ | JOHE4933 | *MAT*α *ago1::NAT* | H99 *ago1*Δ, RNAi mutant | (25) |
| Bt65+*ZNF3* (SJP581) | JOHE22751 | *MAT***a** *ZNF3* | RNAi-repaired, independent from SJP584 | (20) |
| Bt65+*ZNF3-2* (SJP584) | JOHE22754 | *MAT***a** *ZNF3* | RNAi-repaired, independent from SJP581 | (20) |
| Bt65 *ago1*Δ (SJP676) | JOHE22786 | *MAT***a** *ZNF3 ago1::NAT* | Bt65 *ago1*Δ, RNAi mutant | (20) |
| KN99α *frr1*Δ (JHG54) | JOHE23299 | *MAT*α *frr1*Δ*::NAT* | From 2015 Madhani deletion collection | (35) |
| KN99α *msh2*Δ (SJP241) | JOHE22453 | *MAT*α *msh2*Δ*::NAT* | From 2015 Madhani deletion collection | (35) |
| Bt65+*ZNF3 rde4* (JHG1) | JOHE23288 | *MAT***a** *ZNF3 rde4* | Bt65+*ZNF3 rde4*, constructed on SJP581 background | This study |
| Bt65+*ZNF3 znf3-1* (JHG2) | JOHE23289 | *MAT***a** *ZNF3 znf3* | independent from JHG3, constructed on SJP581 background | This study |
| Bt65+*ZNF3 znf3-2* (JHG3) | JOHE23290 | *MAT***a** *ZNF3 znf3* | independent from JHG2, constructed on SJP581 background | This study |
| XL280α (JDW100) | JOHE23300 | *MAT*α | *C. deneoformans* lab strain | (36) |
| XL280α *rdp1*Δ (JDW109) | JOHE23301 | *MAT*α *rdp1::NAT* | XL280α *rdp1*Δ, RNAi mutant | (36) |
| 37-02 (JDW202) | JOHE23302 | *MAT*α | XL280α passaged strain | (33) |
| 37-02 *rdp1*Δ*-1* (JHG9) | JOHE23303 | *MAT*α *rdp1::NAT* | Independent from JHG16, RNAi mutant | This study |
| 37-02 *rdp1*Δ*-2* (JHG16) | JOHE23304 | *MAT*α *rdp1::NAT* | Independent from JHG9, RNAi mutant | This study |
| MA28-H99 30°C (JHG55) | JOHE23291 | *MAT*α | H99 passaged strain | This study |
| MA28-Bt210 30°C (JHG56) | JOHE23292 | *MAT*α | Bt210 passaged strain | This study |

| | | | | |
|---|---|---|---|---|
| MA28-LP-RSA2296 30°C (JHG57) | JOHE23293 | *MAT*α | LP-RSA2296 passaged strain | This study |
| MA28-LP-RSA2296 37°C (JHG58) | JOHE23294 | *MAT*α | LP-RSA2296 passaged strain | This study |
| MA56-H99 30°C (JHG59) | JOHE23295 | *MAT*α | H99 passaged strain | This study |
| MA56-Bt210 30°C (JHG60) | JOHE23296 | *MAT*α | Bt210 passaged strain | This study |
| MA56-LP-RSA2296 30°C (JHG61) | JOHE23297 | *MAT*α | LP-RSA2296 passaged strain | This study |
| MA56-LP-RSA2296 37°C (JHG62) | JOHE23298 | *MAT*α | LP-RSA2296 passaged strain | This study |
| Bt210 F1 progeny P1 (SJP720) | JOHE22924 | *MAT*α *znf3 RDE4* | Bt65 x Bt210 F1 progeny#1 | This study |
| Bt210 F1 progeny P2 (SJP721) | JOHE22925 | *MAT*α *znf3 rde4* | Bt65 x Bt210 F1 progeny#2 | This study |
| Bt210 F1 progeny P3 (SJP722) | JOHE22926 | *MAT*α *znf3 RDE4* | Bt65 x Bt210 F1 progeny#3 | This study |
| Bt210 F1 progeny P4 (SJP723) | JOHE22927 | *MAT*α *znf3 RDE4* | Bt65 x Bt210 F1 progeny#4 | This study |
| Bt210 F1 progeny P5 (SJP724) | JOHE22928 | *MAT***a** *znf3 RDE4* | Bt65 x Bt210 F1 progeny#5 | This study |
| Bt210 F1 progeny P6 (SJP725) | JOHE22929 | *MAT***a** *znf3 RDE4* | Bt65 x Bt210 F1 progeny#6 | This study |
| Bt210 F1 progeny P7 (SJP726) | JOHE22930 | *MAT***a** *ZNF3 RDE4* | Bt65 x Bt210 F1 progeny#7 | This study |
| Bt210 F1 progeny P8 (SJP727) | JOHE22931 | *MAT*α *znf3 RDE4* | Bt65 x Bt210 F1 progeny#8 | This study |
| Bt210 F1 progeny P9 (SJP728) | JOHE22932 | *MAT***a** *ZNF3 RDE4* | Bt65 x Bt210 F1 progeny#9 | This study |
| Bt210 F1 progeny P10 (SJP729) | JOHE22933 | *MAT*α *znf3 RDE4* | Bt65 x Bt210 F1 progeny#10 | This study |
| Bt210 F1 progeny P11 (SJP730) | JOHE22934 | *MAT*α *znf3 rde4* | Bt65 x Bt210 F1 progeny#11, non-hypermutator | This study |
| Bt210 F1 progeny P12 (SJP731) | JOHE22935 | *MAT***a** *ZNF3 rde4* | Bt65 x Bt210 F1 progeny#12, hypermutator | This study |
| Bt210 F1 progeny P13 (SJP732) | JOHE22936 | *MAT*α *ZNF3 RDE4* | Bt65 x Bt210 F1 progeny#13 | This study |
| Bt210 F1 progeny P14 (SJP733) | JOHE22937 | *MAT***a** *ZNF3 RDE4* | Bt65 x Bt210 F1 progeny#14 | This study |
| Bt210 F1 progeny P15 (SJP734) | JOHE22938 | *MAT*α *znf3 rde4* | Bt65 x Bt210 F1 progeny#15 | This study |
| Bt210 F1 progeny P16 (SJP735) | JOHE22939 | *MAT***a** *ZNF3 RDE4* | Bt65 x Bt210 F1 progeny#16 | This study |
| Bt210 F1 progeny P17 (SJP736) | JOHE22940 | *MAT*α *znf3 RDE4* | Bt65 x Bt210 F1 progeny#17 | This study |
| Bt210 F1 progeny P18 (SJP737) | JOHE22941 | *MAT***a** *ZNF3 RDE4* | Bt65 x Bt210 F1 progeny#18 | This study |
| Bt210 F1 progeny P19 (SJP738) | JOHE22942 | *MAT***a** *ZNF3 RDE4* | Bt65 x Bt210 F1 progeny#19 | This study |
| Bt210 F1 progeny P20 (SJP739) | JOHE22943 | *MAT***a** *ZNF3 rde4* | Bt65 x Bt210 F1 progeny#20, non-hypermutator | This study |
| Bt210 F1 progeny P21 (SJP740) | JOHE22944 | *MAT*α *znf3 rde4* | Bt65 x Bt210 F1 progeny#21 | This study |
| Bt210 F1 progeny P22 (SJP741) | JOHE22945 | *MAT*α *ZNF3 rde4* | Bt65 x Bt210 F1 progeny#22, non-hypermutator | This study |
| Bt210 F1 progeny P23 (SJP742) | JOHE22946 | *MAT***a** *znf3 RDE4* | Bt65 x Bt210 F1 progeny#23 | This study |
| Bt210 F1 progeny P24 (SJP743) | JOHE22947 | *MAT*α *ZNF3 RDE4* | Bt65 x Bt210 F1 progeny#24 | This study |
| Bt210 F1 progeny P25 (SJP744) | JOHE22948 | *MAT*α *ZNF3 rde4* | Bt65 x Bt210 F1 progeny#25, hypermutator | This study |
| Bt210 F1 progeny P26 (SJP745) | JOHE22949 | *MAT*α *ZNF3 rde4* | Bt65 x Bt210 F1 progeny#26, non-hypermutator | This study |
| Bt210 F1 progeny P27 (SJP746) | JOHE22950 | *MAT*α *ZNF3 RDE4* | Bt65 x Bt210 F1 progeny#27 | This study |

| | | | | |
|---|---|---|---|---|
| Bt210 F1 progeny P28 (SJP747) | JOHE22951 | *MAT***a** *ZNF3 rde4* | Bt65 x Bt210 F1 progeny#28, non-hypermutator | This study |
| Bt210 F1 progeny P29 (SJP748) | JOHE22952 | *MAT***a** *znf3 RDE4* | Bt65 x Bt210 F1 progeny#29 | This study |
| Bt210 F1 progeny P30 (SJP749) | JOHE22953 | *MAT***a** *ZNF3 rde4* | Bt65 x Bt210 F1 progeny#30, non-hypermutator | This study |
| Bt210 F1 progeny P31 (SJP750) | JOHE22954 | *MAT***a** *znf3 RDE4* | Bt65 x Bt210 F1 progeny#31 | This study |
| Bt210 F1 progeny P32 (SJP751) | JOHE22955 | *MAT***a** *ZNF3 rde4* | Bt65 x Bt210 F1 progeny#32, non-hypermutator | This study |
| Bt210 F1 progeny P33 (SJP752) | JOHE22956 | *MATα ZNF3 RDE4* | Bt65 x Bt210 F1 progeny#33 | This study |
| Bt210 F1 progeny P34 (SJP753) | JOHE22957 | *MAT***a** *ZNF3 rde4* | Bt65 x Bt210 F1 progeny#34, non-hypermutator | This study |
| Bt210 F1 progeny P35 (SJP754) | JOHE22958 | *MAT***a** *znf3 RDE4* | Bt65 x Bt210 F1 progeny#35 | This study |
| Bt210 F1 progeny P36 (SJP755) | JOHE22959 | *MATα znf3 rde4* | Bt65 x Bt210 F1 progeny#36 | This study |
| Bt210 F1 progeny P37 (SJP756) | JOHE22960 | *MAT***a** *znf3 RDE4* | Bt65 x Bt210 F1 progeny#37 | This study |
| Bt210 F1 progeny P38 (SJP757) | JOHE22961 | *MATα znf3/ZNF3 rde4* | Bt65 x Bt210 F1 progeny#38 | This study |
| Bt210 F1 progeny P39 (SJP758) | JOHE22962 | *MATα ZNF3 RDE4* | Bt65 x Bt210 F1 progeny#39 | This study |
| Bt210 F1 progeny P40 (SJP759) | JOHE22963 | *MATα ZNF3 RDE4* | Bt65 x Bt210 F1 progeny#40 | This study |
| Bt210 F1 progeny P41 (SJP760) | JOHE22964 | *MATα ZNF3 RDE4* | Bt65 x Bt210 F1 progeny#41 | This study |
| Bt210 F1 progeny P42 (SJP761) | JOHE22965 | *MATα znf3 RDE4* | Bt65 x Bt210 F1 progeny#42 | This study |
| Bt210 F1 progeny P43 (SJP762) | JOHE22966 | *MAT***a** *ZNF3 rde4* | Bt65 x Bt210 F1 progeny#43, non-hypermutator | This study |
| Bt210 F1 progeny P44 (SJP763) | JOHE22967 | *MAT***a** *znf3 rde4* | Bt65 x Bt210 F1 progeny#44 | This study |
| Bt210 F1 progeny P45 (SJP764) | JOHE22968 | *MATα znf3 RDE4* | Bt65 x Bt210 F1 progeny#45 | This study |
| Bt210 F1 progeny P46 (SJP765) | JOHE22969 | *MATα znf3 rde4* | Bt65 x Bt210 F1 progeny#46 | This study |
| Bt210 F1 progeny P47 (SJP766) | JOHE22970 | *MAT***a** *znf3 rde4* | Bt65 x Bt210 F1 progeny#47 | This study |
| Bt210 F1 progeny P48 (SJP767) | JOHE22971 | *MATα ZNF3 rde4* | Bt65 x Bt210 F1 progeny#48, non-hypermutator | This study |
| Bt210 F1 progeny P49 (SJP768) | JOHE22972 | *MATα znf3 rde4* | Bt65 x Bt210 F1 progeny#49 | This study |
| Bt210 F1 progeny P50 (SJP769) | JOHE22973 | *MATα znf3 RDE4* | Bt65 x Bt210 F1 progeny#50 | This study |
| Bt210 F1 progeny P51 (SJP770) | JOHE22974 | *MAT***a** *znf3 rde4* | Bt65 x Bt210 F1 progeny#51 | This study |
| Bt210 F1 progeny P52 (SJP771) | JOHE22975 | *MAT***a** *znf3 rde4* | Bt65 x Bt210 F1 progeny#52 | This study |
| Bt210 F1 progeny P53 (SJP772) | JOHE22976 | *MAT***a** *ZNF3 RDE4* | Bt65 x Bt210 F1 progeny#53 | This study |
| Bt210 F1 progeny P54 (SJP773) | JOHE22977 | *MAT***a** *ZNF3 RDE4* | Bt65 x Bt210 F1 progeny#54 | This study |
| Bt210 F1 progeny P55 (SJP774) | JOHE22978 | *MAT***a** *znf3 rde4* | Bt65 x Bt210 F1 progeny#55 | This study |
| Bt210 F1 progeny P56 (SJP775) | JOHE22979 | *MAT***a** *znf3 RDE4* | Bt65 x Bt210 F1 progeny#56 | This study |
| Bt210 F1 progeny P57 (SJP776) | JOHE22980 | *MAT***a** *znf3 rde4* | Bt65 x Bt210 F1 progeny#57, hypermutator | This study |
| H99 *crg1*Δ F1 progeny P24 (SJP272) | JOHE22481 | *znf3* | Bt65 x H99 *crg1*Δ F1 progeny#24, hypermutator | (20) |
| H99 *crg1*Δ F1 progeny P34 (SJP282) | JOHE22491 | *znf3* | Bt65 x H99 *crg1*Δ F1 progeny#34, hypermutator | (20) |

**Table S7. Primers used in this study.**

| Primer Name | Primer Sequence (5’-3’) | Purpose |
|---|---|---|
| JOHE52646/JH1 | CTAAGGAATCAATGCCATCC | *CnAGO1* genotyping F |
| JOHE52647/JH2 | TACCCTCGTAAATAACGGAAG | *CnAGO1* genotyping R |
| JOHE52648/JH3 | GTACAAAGGATATGAGGAGGGA | *CnRDE4* genotyping F |
| JOHE52649/JH4 | ATGTGCTGAAGGTAGGCAAT | *CnRDE4* genotyping R |
| JOHE52650/JH5 | GTCTCGTCCTCCTCTTCACC | *CnZNF3* genotyping F |
| JOHE52651/JH6 | CCCATCCTGGCTATCCTCTA | *CnZNF3* genotyping R |
| JOHE52652/JH7 | GGTGTCATGGTCGGTATGGG | *CnACTIN* genotyping F |
| JOHE52653/JH8 | GATACGGAGGATAGCGTGGG | *CnACTIN* genotyping R |
| JOHE52686/JH14 | CGGCTAGTGAAGAACGAACTTG | To amplify *CnZNF3* allele exchange donor F |
| JOHE52687/JH15 | GCATCAAGAAAGCAGCATTTG | To amplify *CnZNF3* allele exchange donor R |
| JOHE52688/JH16 | GTTTCACAGGCTTCTCAGTT | To amplify *CnRDE4* allele exchange donor F |
| JOHE52689/JH17 | TAATCATGTCCTCCGATTGC | To amplify *CnRDE4* allele exchange donor R |
| JOHE52704/JH28 | AAAACGACGGCCAGTGAATT | To amplify 1st part of *NAT* selectable marker F |
| JOHE52705/JH29 | AAGCGACGAATGTTCTGGAGT | To amplify 1st part of *NAT* selectable marker R |
| JOHE52706/JH30 | GAAGAAATGGATGTGCGTTGG | To amplify 2nd part of *NAT* selectable marker F |
| JOHE52707/JH31 | AGCTATGACCATGATTACGCCA | To amplify 2nd part of *NAT* selectable marker R |
| JOHE52712/JH36 | TGTAAAACGACGGCCAGTG | M13F, to amplify Cas9 |
| JOHE52713/JH37 | GCGGATAACAATTTCACACAGG | M13R, to amplify Cas9 |
| JOHE52801/JH46 | CGTGACTACATTTGATTCCGTC | *CnGWC1* genotyping F |
| JOHE52802/JH47 | TTCAAGCGTGATTCAGAGGC | *CnGWC1* genotyping R |
| JOHE52803/JH48 | TTGCCTGCTGAGTTGTTCAT | *CnQIP1* genotyping F |
| JOHE52804/JH49 | CTTCTTCCATTACGCCTTCC | *CnQIP1* genotyping R |
| JOHE52820/JH60 | CTATGTCCGCTCACCGCTAC | *CnFRR1* long amplification F |
| JOHE52821/JH61 | TGCTCGGGAAAGACGAAACT | *CnFRR1* long amplification R |
| JOHE42462/ZC7 | ACAGTCACGAGCCCTGAAAC | *CnFRR1* short amplification F |
| JOHE42463/ZC8 | AGTCGGAGGTTTGGACAGTG | *CnFRR1* short amplification R |
| JOHE52826/JH66 | CTTTCAAATAGGGTATAATTCATCACTATTATTG | *CnRDE4* point mutation F |
| JOHE52827/JH67 | TCAGCGTGTGTTTATGCAG | *CnRDE4* point mutation R |
| JOHE52828/JH68 | TTTGGAGATTAGGAAGCTGAGC | *CnZNF3* point mutation F |
| JOHE52829/JH69 | TCTTCCACCAACAGAGGG | *CnZNF3* point mutation R |

| | | |
|---|---|---|
| JOHE52830/JH70 | AATTGGAGCTCCACCGCG | *CnCas9*_PCnU6/F, gRNA assembly |
| JOHE52831/JH71 | GGGAACAAAAGCTGGTACCG | *CnCas9*_sgRNA/R, gRNA assembly |
| JOHE52832/JH72 | GAGTGCTGTGGTGAAAGAGATGTTTTAGAGCTAGAAATAGCAAGTT | *CnSH1*_sgRNAF, gRNA assembly |
| JOHE52833/JH73 | ATCTCTTTCACCACAGCACTCAACAGTATACCCTGCCGGTG | *CnSH1*_PCnU6-R, gRNA assembly |
| JOHE52834/JH74 | GTTGGTGGAAGATTTGGAGATGTTTTAGAGCTAGAAATAGCAAGTT | *CnZNF3*_exchange_g3_sgRNA/F, gRNA assembly |
| JOHE52835/JH75 | ATCTCCAAATCTTCCACCAACAACAGTATACCCTGCCGGTG | *CnZNF3*_ exchange_g3_PCnU6/R, gRNA assembly |
| JOHE52838/JH78 | GCACACGCTGACTTTCAAATAGTTTTAGAGCTAGAAATAGCAAGTT | *CnRDE4*_ exchange_g2_sgRNAF, gRNA assembly |
| JOHE52839/JH79 | TATTTGAAAGTCAGCGTGTGCAACAGTATACCCTGCCGGTG | *CnRDE4*_ exchange_g2_PCnU6-R, gRNA assembly |
| JOHE52906/JH92 | GAAAGGTCGGCAGTTCACACGTTTTAGAGCTAGAAATAGCAAGTT | *CdRDP1*_g1_sgRNAF, gRNA assembly |
| JOHE52907/JH93 | GTGTGAACTGCCGACCTTTCAACAGTATACCCTGCCGGTG | *CdRDP1*_g1_PCnU6-R, gRNA assembly |
| JOHE52960/JH108 | GAGATGGGGCTGGATGTATGGGTTTTAGAGCTAGAAATAGCAAGTT | *CdRDP1*_g2_sgRNAF, gRNA assembly |
| JOHE52961/JH109 | CCATACATCCAGCCCCATCTcAACAGTATACCCTGCCGGTG | *CdRDP1*_g2_PCnU6-R, gRNA assembly |
| JOHE53079/JH139 | CCCGCTGACTGGGTGAATAG | *CnRDE4*_exchange2_spanning F |
| JOHE53080/JH140 | GACAGACGACTGCGACGACA | *CnRDE4*_exchange2_ spanning R |
| JOHE53085/JH145 | TGTTGTCGCATCCACAGCTAC | *CnZNF3*_exchange3_ spanning F |
| JOHE53086/JH146 | TTGCAGACGGGAAGCTCAATC | *CnZNF3*_exchange3_ spanning R |
| JOHE53091/JH151 | ATCGCTCCATGATGGTGATAC | Cas9_Hm_genotyping F |
| JOHE53092/JH152 | GAGTTCGTCTACGGTGACTACAA | Cas9_Hm_genotyping R |
| JOHE53093/JH153 | GCAAGGACAATGGTAACCTG | sgRNA_HmU6_genotyping F |
| JOHE53094/JH154 | CCTCACTAAAGGGAACAAAA | sgRNA_Hmscaffold_genotyping R |
| JOHE53095/JH155 | AGAGGTAGTAGCGGAGAAGAT | *CnSH1*_genotyping F |
| JOHE53096/JH156 | TGATGAATGGATTAGGGATT | *CnSH1*_ genotyping R |
| JOHE53144/JH170 | CGACATATTGGGTATCCACG | *CdRDP1*_Ko_upstream F |
| JOHE53145/JH171 | AATTCACTGGCCGTCGTTTTGGGTCTATGGGCTATGCTTT | *CdRDP1*_Ko_upstream R |
| JOHE53146/JH172 | TGGCGTAATCATGGTCATAGCTTACCCGTTGCTACGCCCTAC | *CdRDP1*_Ko_downstream F |
| JOHE53147/JH173 | CGCCCATTCACCCACTAAAA | *CdRDP1*_Ko_downstream R |
| JOHE52854/JH80 | AAGTGATCTTGCGGACCTAA | *CdRDP1*_internal F |
| JOHE52855/JH81 | ATGATGGCAGCAACATCTAG | *CdRDP1*_ internal R |
| JOHE53159/JH174 | CTGGTCGATGATATTGTCGTAA | *CdRDP1*_5pjunction F |

| | | |
|---|---|---|
| JOHE53160/JH175 | TCGGAAGACAGTATCGGTTG | *CdRDP1*_5pjunction R |
| JOHE53161/JH176 | CATCCTTGTTGGTGCTGTAA | *CdRDP1*_3pjunction F |
| JOHE53162/JH177 | GAGCCTCAGACTTCTGCTTG | *CdRDP1*_3pjunction R |
| JOHE53311/JH180 | GCAGGGATGGAACATTGATA | *CdFRR1*_long amplification F |
| JOHE53312/JH181 | GAGACCCTCACCTGTTTGCT | *CdFRR1*_long amplification R |
| JOHE52692/JH20 | AAGGACCTCTACGGTAACATTG | *CdACTIN*_genotyping F |
| JOHE52693/JH21 | GACCAGACTCGTCGTACTCG | *CdACTIN*_genotyping R |
| JOHE51977/SJP227 | TTCGTTATGTCCCGCTAAAGG | *CnQIP1* upstream 5' UTR F |
| JOHE51978/SJP228 | GCCTGTGAAAGATGGGTTCTC | *CnQIP1* in-gene (in Intron 2) R |

**Legends for Datasets S1 to S15**

**Dataset S1.** Sanger confirmation for putative RNAi loss-of-function mutations.

**Dataset S2.** Mutation rates, 95% confidence intervals, and raw counts for the fluctuation assay are presented in the left panel of Fig. 2.

**Dataset S3.** Mutation rates, 95% confidence intervals, and raw counts for the fluctuation assay are presented in the right panel of Fig. 2.

**Dataset S4.** Mutation rates, 95% confidence intervals, and raw counts for the fluctuation assay are presented in Fig. 3B.

**Dataset S5.** Raw amplicon sequencing results for the allele exchange strains are presented in Fig. 3C.

**Dataset S6.** Mutation rates, 95% confidence intervals, and raw counts for the fluctuation assay are presented in Fig. S6A.

**Dataset S7.** Raw amplicon sequencing results for the passaged strains are presented in Fig. S6B.

**Dataset S8.** Mutation rates, 95% confidence intervals, and raw counts for the fluctuation assay are presented in Fig. 5B.

**Dataset S9.** Raw amplicon sequencing results for the passaged strains are presented in Fig. 5C.

**Dataset S10.** Contingency table for chi-square test of association between actual and expected inheritance at the mating type, *ZNF3*, and *RDE4* loci in Bt65 x Bt210 F1 progeny.

**Dataset S11.** Mutation rates, 95% confidence intervals, and raw counts for the fluctuation assay are presented in Fig. 6A.

**Dataset S12.** Raw amplicon sequencing results for the F1 progeny are presented in Fig. 6C.

**Dataset S13.** Mutation rates, 95% confidence intervals, and raw counts for the fluctuation assay are presented in Fig. S11.

**Dataset S14.** Predicted repeat contents in Bt210, LP-RSA2296 and H99.

**Dataset S15**. Whole-genome assemblies used for transposon copy number estimation.
